# A human-accelerated enhancer rewires *FOXC1*-dependent skeletal development

**DOI:** 10.64898/2026.11.25.769351

**Authors:** Yue Zhang, Jiamiao Yuan, Xuerui Zeng, Chunmei Li, Yongqin Wang, Jichao Yin, Fengyun Zhang, Bing Su

## Abstract

The evolution of human skeletal morphology involved modifications to conserved developmental programs, but the underlying regulatory mechanisms remain poorly understood. Here we integrate single-cell transcriptomic atlases of developing human, macaque and mouse limbs with comparative chromatin interaction maps of chondrocytes. We identify human-biased transcriptional programs within conserved osteochondral lineages and link divergent chromatin interactions to human-specific sequence changes. These analyses identify zooHAR.30, a human accelerated region that contacts the promoter of *FOXC1*, a key regulator of skeletal development. Human-specific substitutions within zooHAR.30 alter transcription factor binding, and the human sequence exhibits enhanced regulatory activity in chondrocytes and more spatially restricted activity in the developing limb, predominantly in the zeugopod, compared with the chimpanzee ortholog. Replacing the endogenous human sequence with its chimpanzee ortholog reduces *FOXC1* and *GMDS* expression, impairs cell proliferation and osteochondral differentiation, and alters FOXC1 occupancy across skeletal regulatory elements, including a *GDF5* enhancer implicated in knee morphology and osteoarthritis. Our findings reveal how evolutionary changes in a single enhancer can propagate through a conserved developmental regulatory network, providing a mechanistic link between human-specific sequence divergence and the evolution of skeletal developmental programs.

## Introduction

Habitual bipedalism is a defining feature of human evolution^1^, accompanied by marked changes in skeletal morphology, including relatively short upper limbs, elongated lower limbs, a short and broad pelvis, and specialized knee and foot structures^2,3^. Such morphological innovations are thought to arise largely through modifications to conserved developmental programs^4^. Most limb bones develop through endochondral ossification, in which a cartilage template is progressively replaced by bone^5^. Recent work on human pelvic development has shown how changes in the timing and spatial organization of this process contributed to skeletal adaptations for bipedalism^6^. However, the genetic and developmental mechanisms underlying the evolution of human limb morphology remain poorly understood.

Endochondral ossification involves the coordinated proliferation and differentiation of chondrogenic and osteogenic cell populations. Recent single-cell and spatial transcriptomic atlases have resolved the cellular diversity and developmental trajectories of the human embryonic limb and skeleton^7,8^, revealing broad conservation of limb cell identities between humans and mice^8^. However, comparisons between these evolutionarily distant species cannot readily distinguish human-lineage changes from those shared more broadly among primates. Incorporating non-human primates into comparative developmental analyses provides an evolutionary framework for identifying human-biased cellular and transcriptional programs and investigating their contributions to skeletal diversification.

Evolutionary changes in gene regulation are a major source of morphological diversity^4,9^. Within conserved developmental regulatory networks, changes in *cis*-regulatory elements or *trans*-acting factors can modify the timing, spatial distribution and magnitude of gene expression^4,10,11^. Comparative epigenomic studies of embryonic limbs have identified lineage-specific regulatory activities^12^, and recent high-throughput functional assays have revealed extensive *cis*-regulatory divergence in skeletal cell models^13,14^. However, how these sequence-level changes are integrated into the regulatory networks governing skeletal development remains unclear. In particular, the evolutionary conservation and divergence of three-dimensional chromatin interactions linking developmental enhancers to their target genes remain largely unexplored in osteochondral cells.

Human accelerated regions (HARs) are evolutionarily conserved sequences that have undergone accelerated sequence evolution along the human lineage^15–19^, many of which function as developmental enhancers^20^. A notable example is HACNS1, whose human sequence exhibits enhanced activity in the developing forelimb and alters *Gbx2* expression, suggesting a contribution to the evolution of human hand morphology^21,22^. Recent studies have further shown that HARs can act through both conserved and evolutionarily divergent chromatin interactions^19,23^. Yet how human-specific sequence changes within individual enhancers propagate through conserved regulatory networks to alter skeletal developmental programs remains largely unknown.

Here we generated a single-cell transcriptomic atlas of the developing macaque hindlimb and integrated it with human and mouse datasets to identify human-biased transcriptional programs within conserved osteochondral lineages. By combining comparative chromatin interaction maps with developmental transcriptomic and epigenomic profiles, we identified zooHAR.30, a human accelerated enhancer within the *FOXC1–GMDS* regulatory landscape. Human-specific sequence changes at zooHAR.30 alter transcription factor binding, enhance regulatory activity in chondrocytes and modify the spatial distribution of enhancer activity in the developing limb. Replacing the endogenous human sequence with the orthologous chimpanzee sequence reduces *FOXC1* and *GMDS* expression, impairs cell proliferation and osteochondral differentiation, and alters FOXC1 occupancy at downstream skeletal regulatory elements, including the *GDF5* enhancer R4 implicated in knee morphology. Together, our findings reveal how evolutionary changes in a single enhancer can reshape a broader developmental regulatory network, providing a mechanistic framework for understanding the regulatory evolution of human skeletal development.

## Results

### Conserved osteochondral programs are associated with human skeletal proportions

Limb morphology arises through coordinated patterning, differentiation and growth across multiple tissues, and evolutionary changes in these developmental programs underlie limb diversification. To define the cellular basis of this divergence, we compared cellular organization and transcriptional programs across developing human^8^, rhesus macaque and mouse hindlimbs^8,24,25^. We generated single-cell RNA sequencing (scRNA-seq) data from five macaque embryos spanning embryonic days 35, 40 and 60 (E35, E40 and E60), yielding nine libraries (**Supplementary Fig. 1a,b and Supplementary Table 1**). Marker-based annotation resolved nine osteochondral cell populations with stage-specific distributions (**Supplementary Fig. 1c,e and Supplementary Table 2**): early interzone cells (EarlyIZ) were predominantly recovered at E35, mesenchymal condensation cells (MesCond) and interzone cells (InterZone) at E40, and resting and articular chondrocytes alongside osteoblasts at E60 (**Supplementary Fig. 1f**). Integrating these profiles with published human and mouse datasets using SysVI^26^ across one-to-one orthologs yielded a cross-species atlas of 404,708 cells spanning 60 cell types across all major developing limb lineages (**Fig. 1a,b, Supplementary Fig. 2a-c** and **Supplementary Table 3**). Cross-species cell-type mapping demonstrated broad correspondence among human, macaque and mouse populations, supporting the evolutionary conservation of limb cellular identities (**Fig. 1c**).

**Fig. 1.**
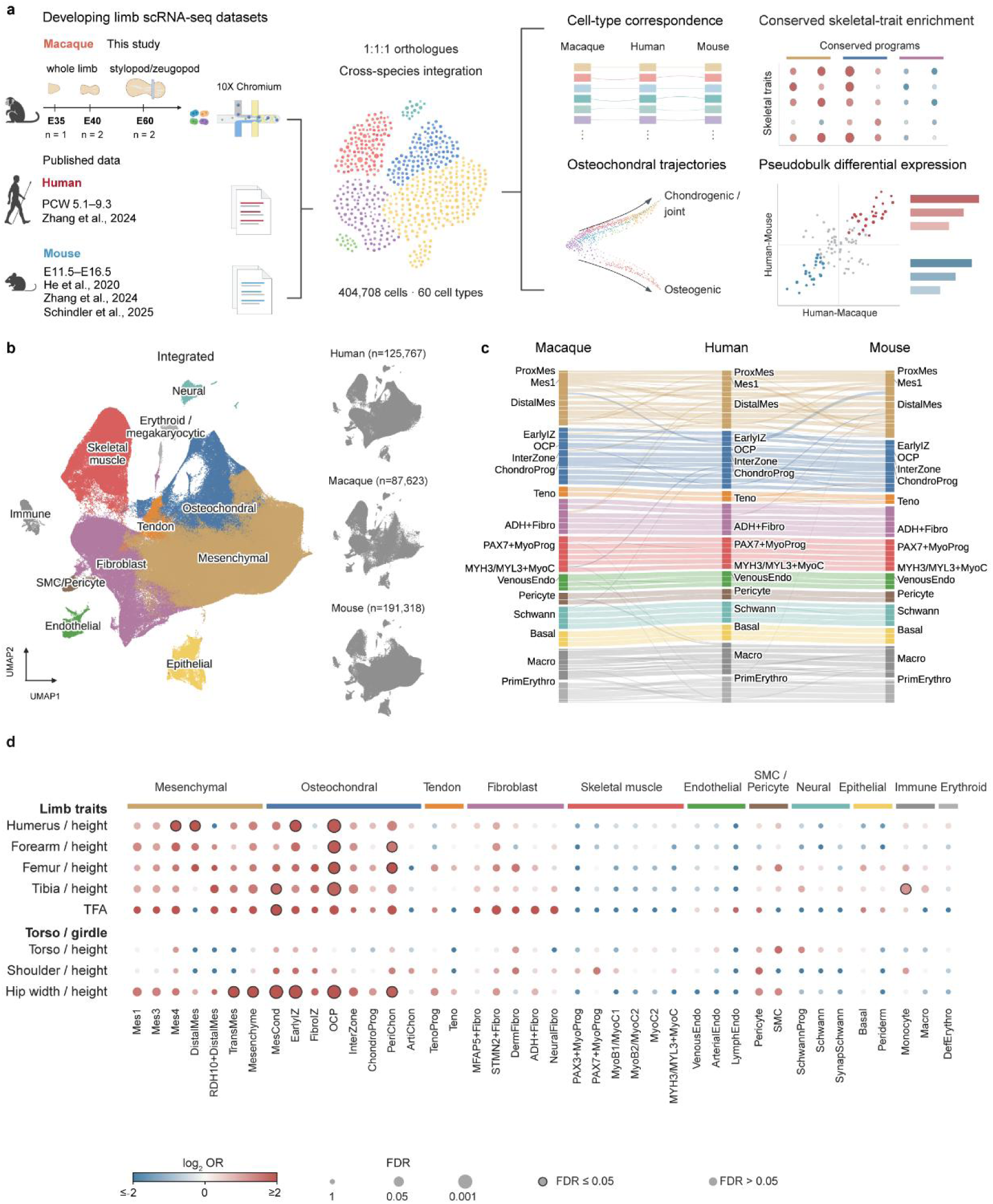
Conserved osteochondral programs are associated with human skeletal proportions. **a**, Study design. Single-cell RNA-seq data generated from macaque limbs at embryonic days (E) 35, E40 and E60 were integrated with published human and mouse limb datasets using one-to-one orthologues. Macaque samples comprised five embryos: one at E35 and two each at E40 and E60. **b**, UMAP of the integrated atlas, colored by major cell lineage. Species-specific views are shown alongside; the number of cells is indicated for each species. **c**, Alluvial plot showing correspondence between annotated cell types across macaque, human and mouse. Colors indicate major lineages. **d**, Enrichment of skeletal-trait-associated genes in conserved cell-type expression programs. Color indicates log₂ odds ratio (OR), and dot size indicates false discovery rate (FDR). Black outlines indicate FDR ≤ 0.05. Enrichment was assessed using one-sided Fisher’s exact tests with Benjamini–Hochberg correction; expressed one-to-one orthologues were used as the background. TFA, tibiofemoral angle.

Because skeletal proportions are governed by the differential growth of individual bones, we next tested whether genes with conserved cell-type-specific expression are enriched for human skeletal proportion traits. Notably, genes associated with limb bone-to-height ratios and hip width-to-height ratios were significantly enriched in several mesenchymal and osteochondral populations, including osteochondral progenitors (OCPs), whereas no such enrichment was observed for axial or pectoral girdle traits, such as torso-to-height and shoulder width-to-height ratios (**Fig. 1d**). Given that limb long bones develop via endochondral ossification, a process in which mesenchymal progenitors differentiate through chondrogenic intermediates prior to bone formation^5^, these findings nominate conserved osteochondral programs as a primary cellular substrate for evolutionary variation in limb proportions, motivating detailed cross-species comparisons of osteochondral development.

### Human-biased transcriptional divergence within conserved osteochondral lineages

To determine how conserved osteochondral cell types give rise to distinct skeletal lineages, we reconstructed their differentiation trajectories and associated transcriptional dynamics across species. Trajectory inference resolved two primary branches corresponding to osteogenic and chondrogenic or joint differentiation (**Fig. 2a** and **Supplementary Fig. 3a,b**). The latter further bifurcated into two sub-trajectories, one extending toward chondrocyte progenitors (ChondroProg) and resting chondrocytes (RestingChon), and the other toward perichondrial cells (PeriChon) and articular chondrocytes (ArtiChon) (Supplementary Fig. 3c–g). OCPs occupied early positions along both major branches, consistent with their bipotent osteogenic and chondrogenic potential (**Fig. 2a,b**). Along these trajectories, OCP-associated genes, including *POMC*, *GMDS* and *FOXC1*, exhibited distinct expression dynamics (**Fig. 2b** and **Supplementary Fig. 3e-g**). Resolving these patterns by species revealed higher expression of *POMC* and *GMDS* in humans at early pseudotime, followed by progressive downregulation during differentiation (**Fig. 2c,d**). By contrast, *COL9A2* and *ACAN*, encoding cartilage extracellular matrix components, peaked later along the chondrogenic trajectory and showed higher fitted expression in humans (**Fig. 2e,f**).

**Fig. 2.**
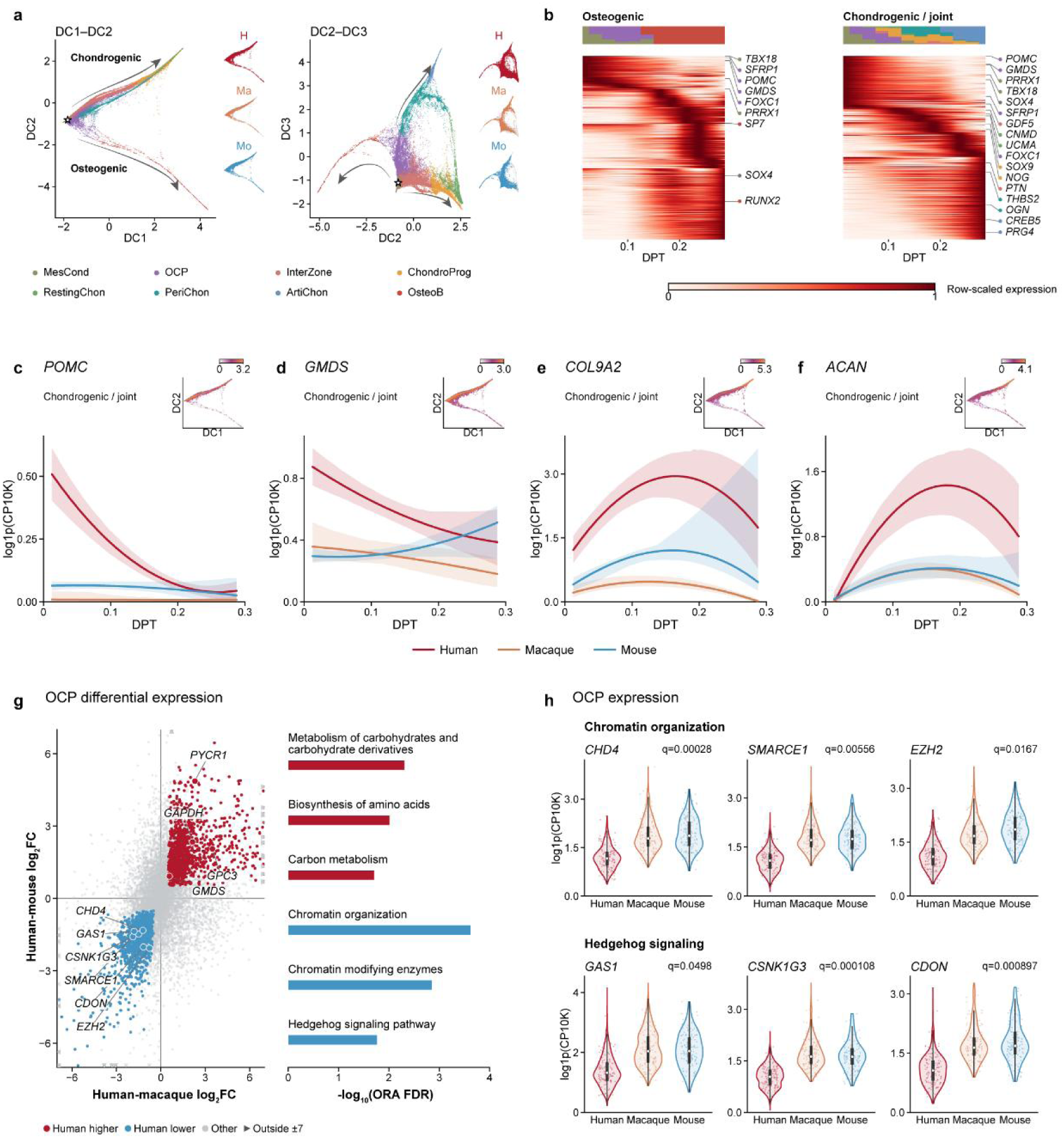
Human-biased transcriptional divergence within conserved osteochondral lineages. **a**, Diffusion map projections of integrated osteochondral cells onto diffusion components (DCs) 1-2 and 2-3. Cells are colored by annotated cell type; species-specific views are shown alongside. Stars indicate the root cell, and arrows indicate inferred trajectory directions. **b**, Gene-expression dynamics along the osteogenic and chondrogenic/joint-associated trajectories. Expression was averaged across samples within each species and then across represented species, smoothed and scaled from 0 to 1 for each gene. Upper bars indicate cell-type composition along diffusion pseudotime (DPT). **c–f**, Expression of POMC (c), GMDS (d), COL9A2 (e) and ACAN (f) along the chondrogenic/joint-associated trajectory. Lines show fitted expression trends; shading indicates exploratory 95% sample-ID bootstrap intervals. Insets show expression projected onto the DC1–DC2 embedding. **g**, Pseudobulk differential expression in osteochondral progenitors (OCPs). Axes show human–macaque and human–mouse log₂ fold changes. Red and blue indicate genes expressed at higher and lower levels in human, respectively; triangles denote values outside the displayed range. Classification required concordant effects in both comparisons, |log₂ fold change| ≥ 0.5 in each and joint FDR ≤ 0.05. Bars show selected over-represented pathways, assessed using one-sided hypergeometric tests with Benjamini–Hochberg correction. **h**, Expression of selected chromatin-organization and Hedgehog-signaling genes in OCPs. Violins show cells with detected expression; overlaid dots are a subsample of cells. White points indicate medians, boxes indicate interquartile ranges (IQRs), and whiskers extend to observations within 1.5 × IQR. q values are from the pseudobulk joint analysis.

To compare OCP transcriptional programs at equivalent developmental stages, we built an age-reference regression model incorporating cell-type composition from human embryonic limb transcriptomes. This model aligned macaque E40 samples with human post-conception week 6 (PCW6), establishing a quantitative framework for cross-species stage matching (**Supplementary Fig. 3h**). Pseudobulk differential expression analysis at matched stages identified concordant transcriptional changes in human OCPs relative to both macaque and mouse counterparts (**Supplementary Table 4**). Genes downregulated in human OCPs were enriched for chromatin organization, chromatin-modifying enzymes and Hedgehog signaling, whereas upregulated genes were enriched for carbohydrate and amino acid metabolism (**Fig. 2g**). Downregulated genes included the chromatin regulators *CHD4*, *SMARCE1* and *EZH2*, alongside the Hedgehog pathway components *CSNK1G3* and *CDON* (**Fig. 2h**).

Several expression shifts observed in OCPs were shared across other early osteochondral populations. Specifically, elevated expression of *SOX4* and *GMDS* in human OCPs extended to InterZone and ChondroProg cells, whereas reduced expression of *BMPR2* was shared with EarlyIZ and InterZone cells, and reduced *TGFBR1* expression was shared with InterZone and ChondroProg cells (**Supplementary Fig. 3i**). Together, these findings reveal human-biased transcriptional divergence in OCPs and related early osteochondral populations within otherwise conserved differentiation trajectories, highlighted by the remodeling of chromatin regulators and raising the question of how *cis*-regulatory landscapes shape these species-specific expression programs.

### Human-gained chromatin interactions preferentially involve enhancers

To investigate the *cis*-regulatory mechanisms underlying the human-biased transcriptional programs observed in early osteochondral progenitors, we examined three-dimensional chromatin interactions across human, macaque and mouse chondrocyte models. Specifically, we profiled the human chondrocyte line CHON-002, the mouse chondrocyte line iMAC and an immortalized macaque chondrocyte line that we established from E60 hindlimb long-bone cartilage via hTERT expression (**Fig. 3a**). Transcriptomic comparison with our developmental single-cell atlas confirmed that all three lines closely resembled early osteochondral progenitor states in vivo (**Supplementary Fig. 4e**). We performed H3K4me3 and H3K27ac HiChIP to map chromatin contacts anchored at active promoters and enhancers across the three species, obtaining highly reproducible contact maps among biological replicates (**Supplementary Fig. 4a**). Multidimensional scaling revealed tight cross-species clustering of H3K4me3 HiChIP libraries, whereas H3K27ac HiChIP libraries exhibited greater evolutionary divergence (**Supplementary Fig. 4b**). Calling interactions independently within each species and comparing contacts across orthologous genomic intervals identified conserved, primate-shared and species-specific loops, including human-gained and human-lost interactions (**Fig. 3b, Supplementary Fig. 4c** and **Supplementary Table 5**).

**Fig. 3.**
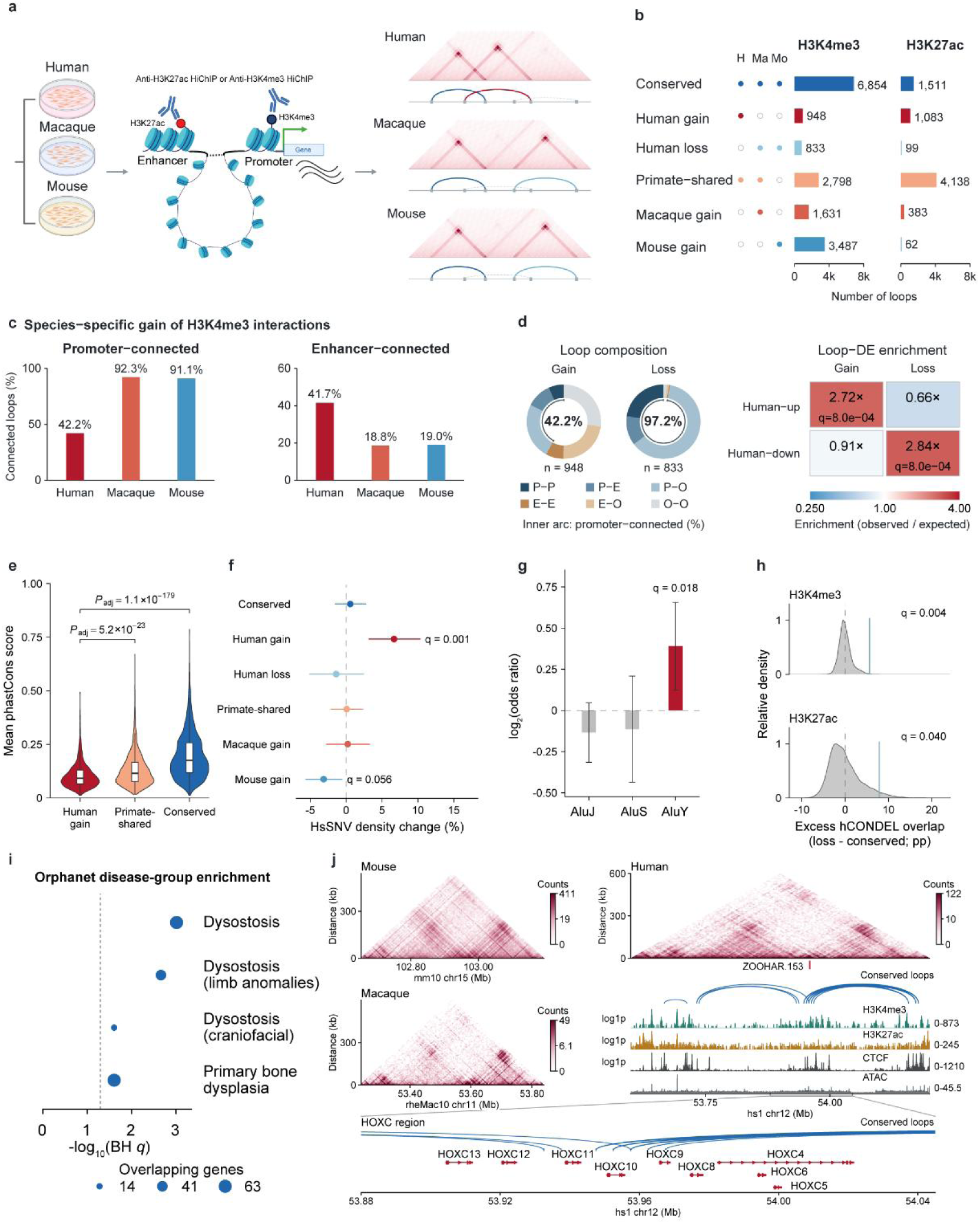
Human-gained chromatin interactions preferentially involve enhancers. **a**, Overview of H3K4me3 and H3K27ac HiChIP in human, macaque and mouse chondrocyte cell lines. **b**, Numbers of loops in six cross-species interaction categories. Filled and open circles indicate detected and undetected interactions, respectively. **c**, Proportions of species-gained H3K4me3 loops connected to promoters or enhancers; the categories can overlap. **d**, Composition of human-gained and human-lost H3K4me3 loops and enrichment for genes deferentially expressed in human. P, promoter; E, enhancer; O, other. Inner arcs indicate the fraction of promoter-connected loops. Heatmap values are observed-to-expected ratios from 10,000 permutations; q values were calculated using two-sided permutation tests and Benjamini-Hochberg correction. **e**, Mean phastCons100way scores of human-gained, primate-shared and conserved loops. Boxes indicate medians and interquartile ranges (IQRs); whiskers extend to observations within 1.5 × IQR. Adjusted P values were calculated using two-sided Wilcoxon rank-sum tests with Benjamini-Hochberg correction. **f**, Change in human-specific single-nucleotide variant density at H3K4me3 loop anchors for each category relative to the other categories. Points and error bars indicate estimates and 95% confidence intervals (CIs) from negative-binomial regression; q values are from two-sided Wald tests with Benjamini-Hochberg correction. **g**, Alu-subfamily enrichment at human-gained versus primate-shared anchors. Bars show log₂ odds ratios; error bars indicate chromosome-level jackknife 95% CIs. q values were adjusted across the three Alu subfamilies. **h**, Matched-permutation analysis of human-specific deletion of conserved sequence (hCONDEL) overlap in human-lost versus conserved loops. Grey curves show centred null distributions; blue lines indicate observed differences. q values were calculated using one-sided permutation tests with Benjamini-Hochberg correction. **i**, Orphanet disease-group enrichment among genes associated with conserved interactions. Dot size indicates overlapping genes; the dashed line marks false discovery rate = 0.05. j, H3K4me3 HiChIP contact maps at the orthologous HOXC region, with human chromatin tracks and conserved interactions.

Analysis of the regulatory architecture of species-specific H3K4me3-anchored contacts revealed a striking shift in human chromatin connectivity. Whereas 92.3% of macaque-gained and 91.1% of mouse-gained interactions were promoter-connected, the corresponding fraction was 42.2% for human-gained interactions. Enhancer-connected interactions accounted for 41.7% of human-gained contacts, compared with 18.8% and 19.0% in macaque and mouse, respectively (**Fig. 3c**). Conversely, 97.2% of human-lost interactions were promoter-connected (**Fig. 3d**). Together, this selective loss of promoter-connected loops alongside the preferential gain of enhancer-connected contacts suggests that evolutionary rewiring of chromatin architecture in humans attenuated canonical promoter–promoter coupling while expanding enhancer-mediated regulation of developmental gene expression. Consistent with functional regulatory coupling, human-gained interactions were significantly enriched at genes upregulated in human chondrocytes relative to macaque and mouse cells, whereas human-lost interactions were enriched at downregulated genes, showing 2.72-fold and 2.84-fold enrichment, respectively (both *q* = 8.0 × 10⁻⁴; **Fig. 3d** and **Supplementary Table 6**). Together, these results demonstrate widespread, evolutionarily driven rewiring of enhancer-promoter connectivity in human chondrocytes that directly aligns with species-specific transcriptional divergence.

### Distinct evolutionary signatures of conserved and divergent chromatin interactions

These evolutionary shifts in chromatin connectivity were accompanied by distinct genomic sequence signatures. Human-gained interactions exhibited lower sequence conservation than both primate-shared and three-species-conserved interactions (**Fig. 3e**), and their anchors were significantly enriched for human-specific single-nucleotide variants relative to all other interaction categories combined (*q* = 0.001; **Fig. 3f**). In addition, the evolutionarily young *AluY* retrotransposon subfamily was enriched at human-gained interactions compared with primate-shared interactions (**Fig. 3g**). Conversely, the anchors of human-lost interactions showed significantly greater overlap with human-specific deletions of conserved sequences (hCONDELs^27^) than those of conserved interactions, a pattern observed for both H3K4me3- and H3K27ac-anchored contacts (**Fig. 3h**). Together, these observations indicate that human-specific chromatin rewiring is closely coupled to recent sequence divergence, including single-nucleotide substitutions, *AluY* insertions and lineage-specific deletions of conserved regulatory elements.

In contrast to these rapidly evolving contacts, interactions conserved across all three species were enriched near genes implicated in rare skeletal developmental disorders, including dysostosis, limb anomalies, craniofacial anomalies and primary bone dysplasia (**Fig. 3i**). Consistent with the established role of CTCF in partitioning *HOX* regulatory domains^28,29^, conserved interactions displayed strong enrichment for CTCF binding (Supplementary Fig. 4d). Given the critical role of *HOXC* genes in hindlimb patterning and development^30^, we examined the *HOXC* cluster in greater detail. Cross-species contact maps revealed a highly conserved interaction architecture across humans, macaques and mice, characterized by invariant loops spanning the cluster and anchored at CTCF-occupied sites (**Fig. 3j**). The *HOXC* locus thus exemplifies an evolutionarily constrained three-dimensional chromatin architecture that safeguards the coordinated spatiotemporal regulation of core limb developmental genes.

### zooHAR.30 exhibits enhanced regulatory activity within the conserved *FOXC1*-*GMDS* landscape

To determine whether sequence divergence contributes to human-specific osteochondral regulation, we integrated cross-species chromatin interactions with developmental transcriptomic and epigenomic profiles to prioritize human accelerated regions (HARs). This analysis highlighted zooHAR.30, an intronic element within *GMDS* that contacts the neighboring *FOXC1* locus (**Fig. 4a**), which encodes a transcription factor essential for both endochondral and intramembranous ossification^31–33^. The zooHAR.30 interval was marked by H3K27ac enrichment and connected to the *FOXC1* promoter via a conserved enhancer-promoter interaction, alongside human-gained enhancer–enhancer contacts across the surrounding locus, situating zooHAR.30 within a regulatory domain combining conserved and evolutionarily divergent connectivity (**Fig. 4a** and **Supplementary Table 5**).

**Fig. 4.**
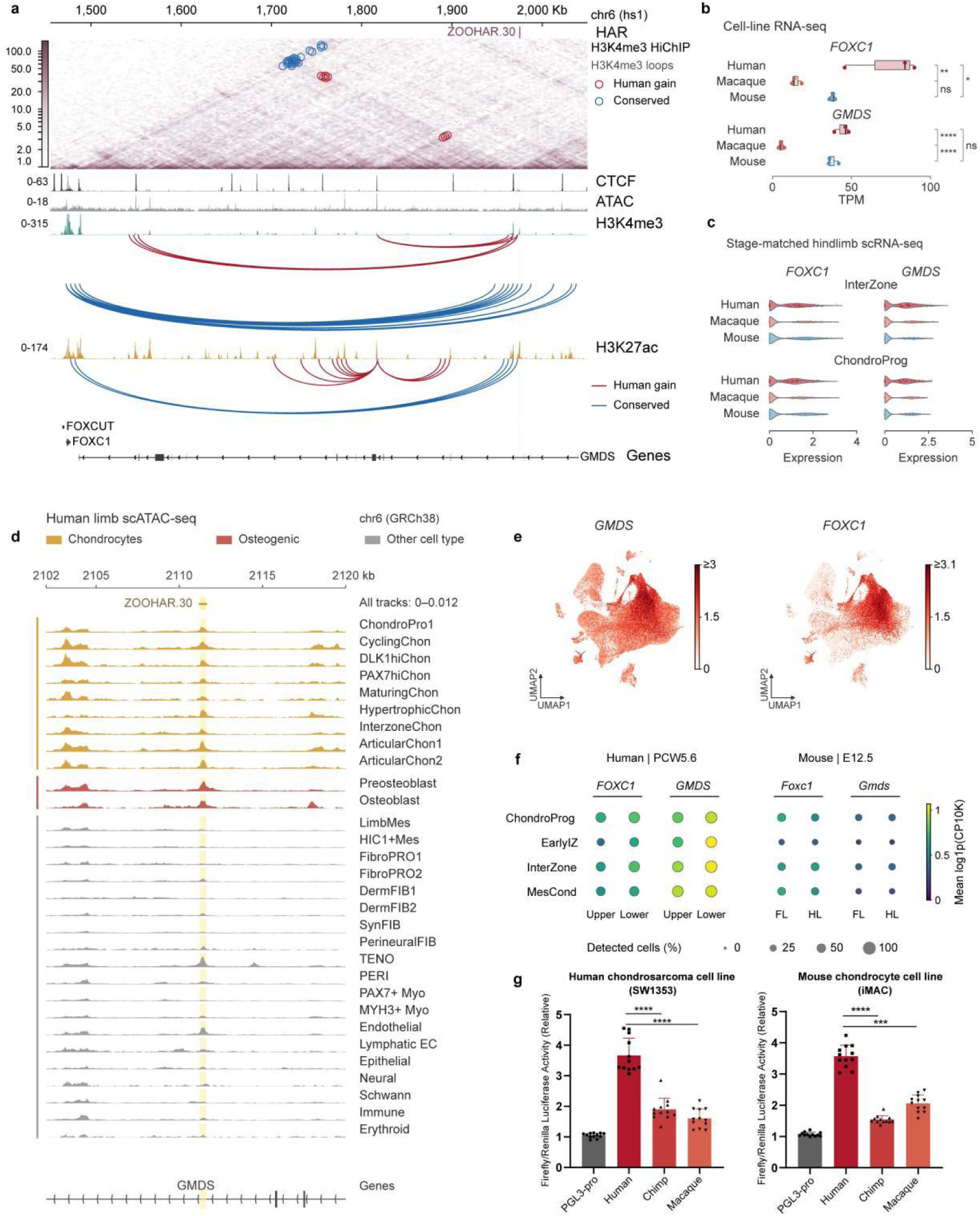
zooHAR.30 exhibits enhanced regulatory activity within the conserved *FOXC1*-*GMDS* landscape. **a**, Chromatin landscape of the zooHAR.30–*FOXC1*/*GMDS* locus in human chondrocytes. The H3K4me3 HiChIP contact map is shown with CTCF, ATAC-seq, H3K4me3 and H3K27ac tracks and gene annotations. Red and blue circles and arcs indicate human-gained and conserved interactions, respectively. Coordinates are in T2T-CHM13/hs1. **b**, RNA-seq expression of FOXC1 and GMDS in human, macaque and mouse chondrocyte cell lines, shown as transcripts per million (TPM; n = 3 samples per species). Boxes indicate medians and interquartile ranges (IQRs); whiskers extend to observations within 1.5 × IQR. Comparisons used one-way ANOVA followed by Tukey’s multiple-comparison test. **c,** Single-cell expression of *FOXC1* and *GMDS* in InterZone and chondroprogenitor (ChondroProg) cells from stage-matched hindlimb samples. Expression is shown as log1p-transformed counts per 10,000 (CP10K). Violin densities include zero-expression cells; dots show a subset of cells. **d,** Cell-type-resolved human limb chromatin accessibility at zooHAR.30. Yellow shading marks the region of interest; tracks share a common signal scale. Coordinates are in GRCh38. **e**, *GMDS* and *FOXC1* expression projected onto the integrated single-cell transcriptomic UMAP. **f**, Upper-lower limb expression of *FOXC1* and *GMDS* and their mouse (forelimb-hindlimb) orthologs in ChondroProg, EarlyIZ, InterZone and MesCond cells from human PCW5.6 and mouse E12.5 samples. Dot color indicates mean log1p(CP10K); dot size indicates the percentage of expressing cells. Human values were averaged equally across matched limb regions, and mouse values across libraries. FL, forelimb; HL, hindlimb. **g**, Dual-luciferase activities of human, chimpanzee and macaque zooHAR.30 sequences in SW1353 and iMAC cells. Firefly activity was normalized to Renilla activity, with pGL3-Promoter as the vector control. Dots indicate individual measurements. The assays were performed in three independent experiments. The human sequence showed higher reporter activity than the chimpanzee and macaque sequences in both cell lines. *, P < 0.05; **, P < 0.01; ***, P < 0.001; ****, P < 0.0001; ns, not significant.

Expression profiling of *FOXC1* and *GMDS* revealed human-biased regulation across both in vitro and developing limb contexts. In chondrocyte lines, *FOXC1* was higher in human than in macaque and mouse cells, whereas *GMDS* exceeded macaque levels but was comparable to mouse (**Fig. 4b**). In vivo, stage-matched hindlimb comparisons showed elevated *GMDS* expression in human InterZone and ChondroProg cells relative to both non-human species across pseudobulk and single-cell profiles (**Fig. 4c** and **Supplementary Table 4**). Across the integrated atlas and spatial transcriptomics, both transcripts overlapped across osteochondral states and localized to human hindlimb skeletal structures (**Fig. 4e** and **Supplementary Fig. 5d**). Within human PCW5.6 limbs, *FOXC1* and *GMDS* also showed consistently higher mean expression in lower than in upper limbs across ChondroProg, EarlyIZ, InterZone and MesCond populations, while such difference was absent in mouse E12.5 limbs (**Fig. 4f**).

Developmental chromatin accessibility and reporter assays further supported a functional regulatory role for zooHAR.30. In human single-cell multi-omic datasets^7^, zooHAR.30 was accessible across chondrogenic and osteogenic populations of the developing limb (**Fig. 4d**), as well as in osteochondral lineages of the developing pelvis and cranial structures undergoing endochondral or intramembranous ossification (**Supplementary Fig. 5a,b**). In luciferase reporter assays comparing the human, chimpanzee and macaque orthologues, the human sequence drove significantly higher transcriptional activity than either non-human primate allele in both human SW1353 and mouse iMAC cells (**Fig. 4g**), demonstrating sequence-dependent gain of enhancer activity across distinct cellular contexts.

We also identified a second candidate, zooHAR.297, near the *ALCAM* promoter within a conserved, CTCF-occupied interaction, where *ALCAM* expression and zooHAR.297 accessibility overlapped in developing limb perichondrial cells^34^ and cranial osteochondral progenitors (**Supplementary Fig. 6a-f**). Together, these findings identify HARs embedded within conserved chromatin interaction landscapes at key skeletal developmental genes, and establish zooHAR.30 as a prime candidate driving human-specific regulatory divergence at the *FOXC1-GMDS* locus.

### The human zooHAR.30 enhancer promotes osteochondral developmental programs

To evaluate the functional necessity of the human zooHAR.30 sequence at its native genomic locus, we used CRISPR-Cas9 genome editing in CHON-002 cells to replace the endogenous human element with the orthologous chimpanzee allele, generating a knock-in line designated Chimp-HAR^KI^ alongside a non-targeting control line (**Fig. 5a**). H3K27ac CUT&Tag profiling revealed reduced acetylation across both zooHAR.30 and the *FOXC1* locus in Chimp-HAR^KI^ cells (**Fig. 5b**), consistent with the attenuated enhancer activity of the chimpanzee allele observed in reporter assays. Sequence replacement also reduced the expression of both *GMDS* and *FOXC1* (**Supplementary Fig. 7d,e**), linking the local chromatin changes to altered transcription at the locus.

**Fig. 5.**
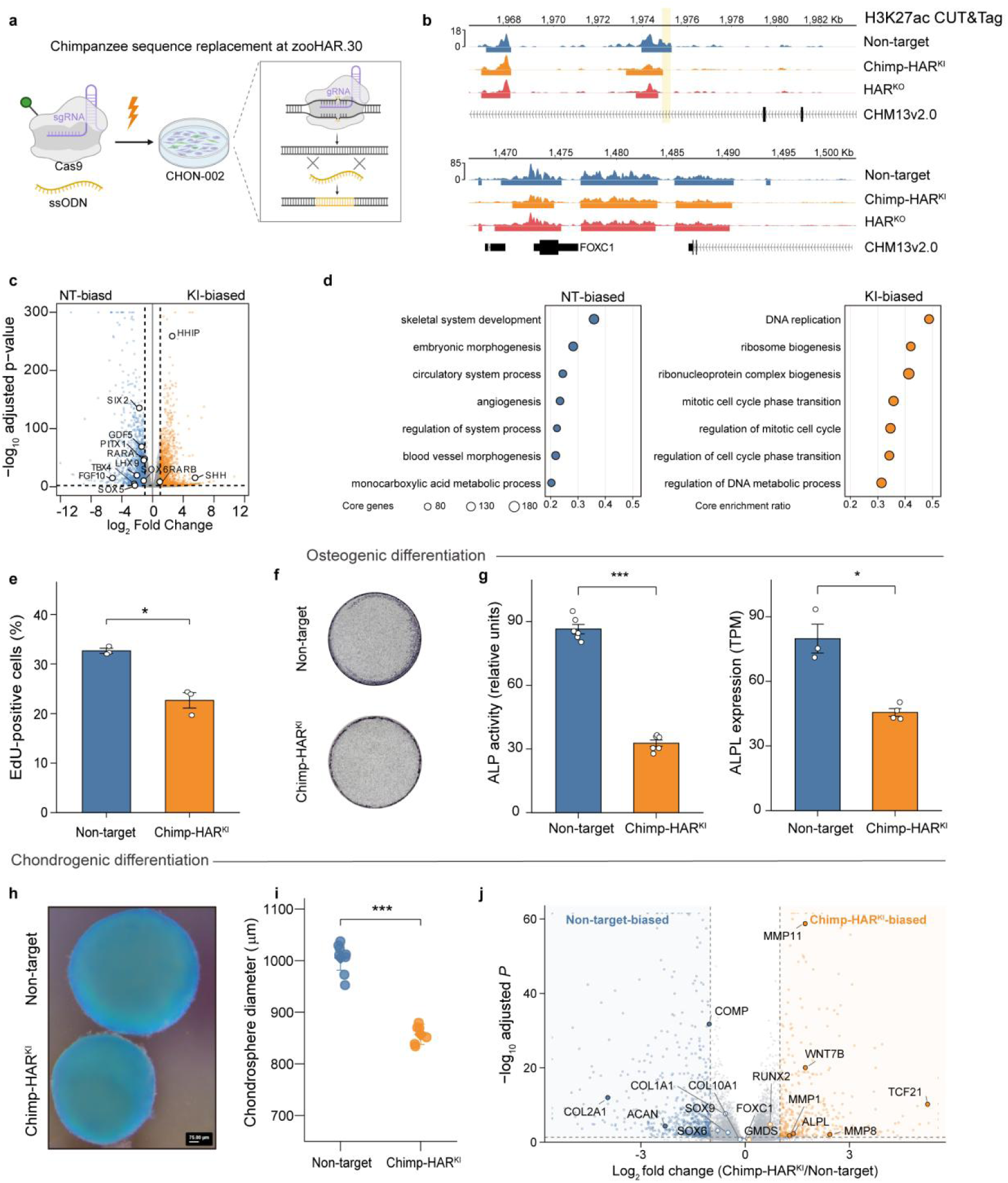
The human zooHAR.30 enhancer promotes osteochondral developmental programs. **a**, CRISPR–Cas9-mediated replacement of the endogenous human zooHAR.30 sequence with the chimpanzee orthologue in CHON-002 cells. **b**, H3K27ac CUT&Tag profiles at zooHAR.30 and the FOXC1 locus in non-targeting control, Chimp-HAR^KI^ and HAR^KO^ cells. Shading marks zooHAR.30. **c**, Differential expression in undifferentiated Chimp-HAR^KI^ versus non-target cells (n = 4 and 3 RNA-seq samples, respectively). Blue and orange indicate higher expression in non-target and Chimp-HAR^KI^ cells, respectively; Benjamini–Hochberg-adjusted P < 0.01 and |log₂ fold change| ≥ 1. **d**, Gene set enrichment analysis of Gene Ontology biological-process terms. The horizontal axis indicates the fraction of each gene set contributing to the core enrichment; dot size indicates the number of core genes. **e**, Percentage of EdU-positive cells measured by flow cytometry (n = 3 independently cultured samples per group). **f**, Representative alkaline phosphatase (ALP) staining after 10 days of osteogenic induction. **g**, ALP activity after 10 days of osteogenic induction and ALPL RNA-seq expression in osteogenically induced cells. ALP activity was measured in six wells per group; ALPL expression was measured in three non-target and four Chimp-HAR^KI^ cell clones. **h**, Representative Alcian blue-stained chondrospheres after 21 days of chondrogenic induction at 5% O₂. Scale bar, 75 µm. **i**, Chondrosphere diameters (n = 8 non-target and 6 Chimp-HAR^KI^ spheroids). **j**, Differential expression in Chimp-HAR^KI^ versus non-target chondrospheres; Benjamini–Hochberg-adjusted P < 0.05 and |log₂ fold change| ≥ 1. Bars and error bars in e,g indicate mean ± s.e.m.; points and error bars in i indicate mean and 95% confidence interval. ALP activity and diameter data are from one of three independent experiments. Pairwise comparisons used two-sided Welch’s t-tests: P = 0.0150 (e), 1.19 × 10⁻⁸ and 0.0293 (g, left and right), and 7.88 × 10⁻⁸ (i). *, P < 0.05; ***, P < 0.001.

Transcriptomic profiling of control and Chimp-HAR^KI^ cells revealed widespread gene expression remodeling following sequence replacement (**Fig. 5c** and **Supplementary Table 7**). Genes downregulated in Chimp-HAR^KI^ cells were enriched for skeletal system development and embryonic morphogenesis, whereas upregulated genes were enriched for DNA replication and cell-cycle processes (**Fig. 5d**). Among the downregulated developmental regulators, *PITX1*, *TBX4* and *FGF10*, which govern hindlimb specification and outgrowth^35–37^, showed concordant reductions in transcript abundance and nearby H3K27ac enrichment (**Fig. 5c** and **Supplementary Fig. 7b**). Sequence replacement also perturbed multiple components of the Hedgehog signaling pathway, increasing *SHH* and *HHIP* expression while reducing *PTCH1* and *GLI3* expression (**Fig. 5c**). Notably, upregulation of *SHH*, a key regulator of limb patterning and growth^38,39^, was accompanied by increased H3K27ac deposition upstream of the locus (**Supplementary Fig. 7c**), mirroring the human-specific downregulation of Hedgehog pathway genes observed in OCPs in vivo.

We next assessed whether these transcriptional and epigenomic shifts affected cellular proliferation, survival and differentiation potential. Chimp-HAR^KI^ cells exhibited reduced proliferation without detectable changes in apoptosis (**Fig. 5e** and **Supplementary Fig. 7f,g**). Upon osteogenic induction, Chimp-HAR^KI^ cells displayed diminished alkaline phosphatase activity and lower expression of the osteogenic marker *ALPL* (**Fig. 5f,g**). Similarly, under three-dimensional chondrogenic culture conditions, Chimp-HAR^KI^ cells formed significantly smaller chondrospheres than non-targeting control cells (Fig. 5h,i and Supplementary Fig. 7h), accompanied by reduced expression of genes encoding cartilage extracellular matrix proteins, including *COMP*, *COL2A1* and *ACAN* (**Fig. 5j** and **Supplementary Table 7**). Together, these endogenous sequence-replacement experiments reveal that human-specific sequence changes at zooHAR.30 promote osteochondral developmental programs by sustaining skeletal gene expression, cellular proliferation and both osteogenic and chondrogenic differentiation potential.

### zooHAR.30 sustains FOXC1 occupancy at skeletal regulatory elements

To determine whether zooHAR.30 sequence replacement also affects the binding landscape of the transcription factor FOXC1, we performed FOXC1 CUT&Tag in non-targeting control and Chimp-HAR^KI^ cells. FOXC1 occupancy decreased at 290 regions but increased at only 15, indicating that binding changes were predominantly reductions (**Fig. 6a** and **Supplementary Table 8**). Regions with decreased binding included loci associated with growth plate signaling and chondrocyte maturation, such as *BMP6* and *PANX3*^40,41^; *HDAC5*, which contributes to the restraint of chondrocyte hypertrophy^42^; and *TGFBI*, which is associated with cartilage matrix maintenance^43^ (**Supplementary Fig. 8a-c**). Reduced FOXC1 binding was also observed at multiple sites upstream of *SHH* (Supplementary Fig. 8d), whose expression increased following sequence replacement (**Fig. 5c**).

**Fig. 6.**
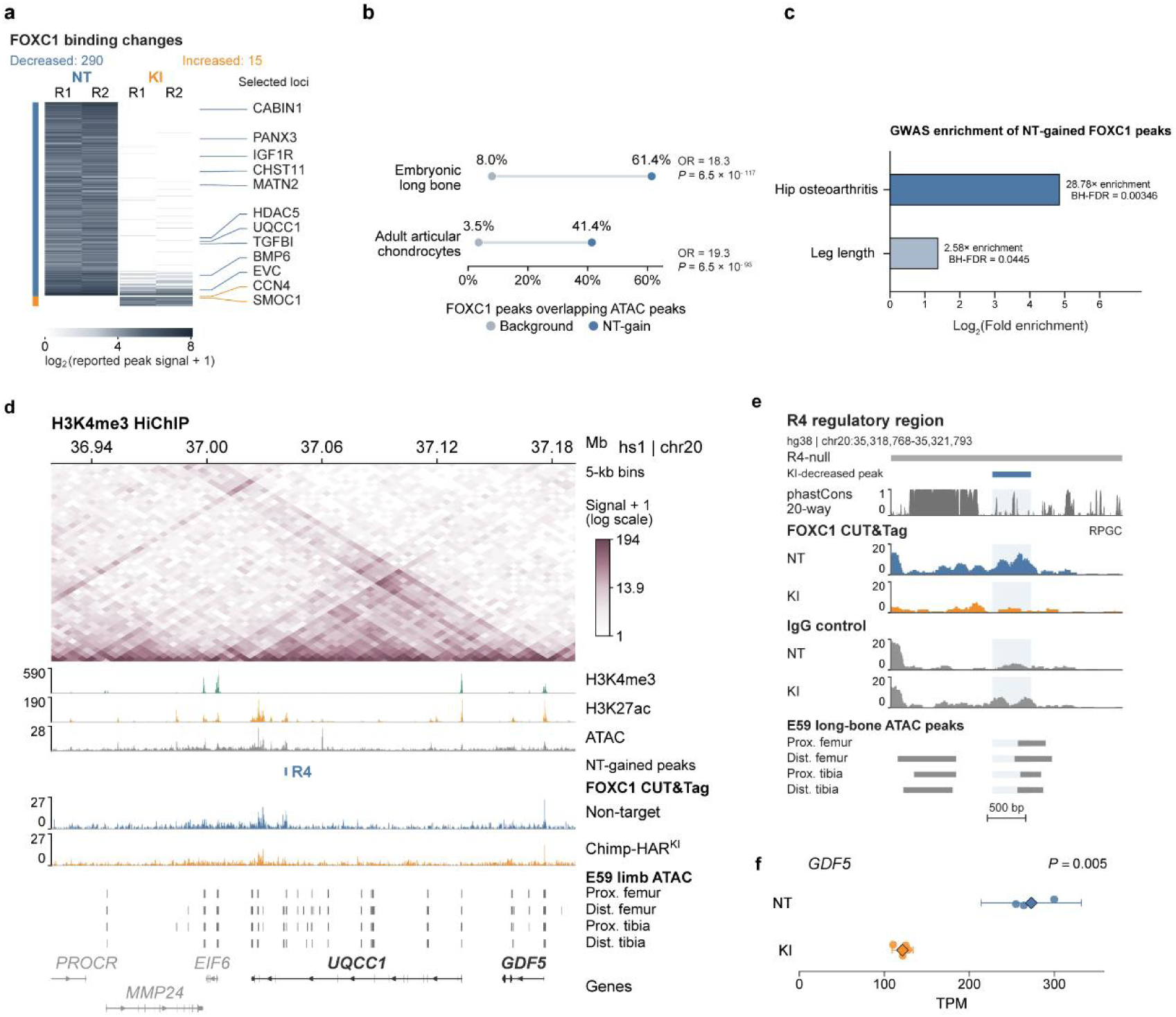
zooHAR.30 sustains FOXC1 occupancy at skeletal regulatory elements. **a**, Differential FOXC1 CUT&Tag signals in non-targeting control (NT) and Chimp-HAR^KI^ (KI) CHON-002 cells, with two replicates per condition. Differential binding was assessed using DiffBind/DESeq2; FDR < 0.05 and |log₂ fold change| ≥ 1. Regions with decreased binding in KI are referred to as NT-gained peaks. **b**, Overlap of NT-gained FOXC1 peaks with accessible chromatin in human embryonic long bone and articular chondrocytes. Points indicate overlap percentages; ORs and two-sided Fisher’s exact test P values are shown. **c**, Enrichment of NT-gained FOXC1 peaks overlapping credible-set variants for hip osteoarthritis and leg length. Bars show log₂ observed-to-expected peak counts; one-sided empirical P values were obtained from 100,000 matched-background simulations and adjusted across the two traits using the Benjamini–Hochberg procedure. **d**, H3K4me3 HiChIP contact map and chromatin tracks at the UQCC1–GDF5 locus in hs1 coordinates. The R4 regulatory region and NT-gained FOXC1 peaks are indicated. **e**, View of the R4 region in hg38 coordinates, showing the previously defined R4-null interval, a KI-decreased FOXC1 peak, phastCons conservation, FOXC1 CUT&Tag and IgG-control signals, and E59 long-bone ATAC peaks. Scale bar, 500 bp. f, RNA-seq expression of GDF5 in NT and KI cells. Circles represent individual samples (n = 3 NT and 4 KI); diamonds and error bars indicate mean and 95% CI. P = 0.0051, two-sided Welch’s t-test.

We compared the FOXC1 peaks reduced in Chimp-HAR^KI^ cells with accessible chromatin regions in skeletal tissues. Of these peaks, 61.4% overlapped accessible regions in human embryonic long bones^44^, compared with 8.0% of background regions; they were also enriched in accessible chromatin from adult articular chondrocytes^45^ (**Fig. 6b**). The same peaks were enriched for variants associated with hip osteoarthritis^46^ and leg length^47^ (**Fig. 6c**). Among them, a FOXC1 peak overlapped R4, a regulatory element at the *UQCC1–GDF5* locus previously shown to influence *GDF5* expression, knee morphology and osteoarthritis risk^44,48,49^(**Fig. 6d,e**). The FOXC1 peak within R4 also overlapped accessible chromatin in developing human long bones, and H3K4me3 HiChIP showed chromatin contacts between the R4-containing region and the *GDF5* locus (**Fig. 6d,e**). Consistent with these changes, *GDF5* expression was reduced in Chimp-HAR^KI^ cells (**Fig. 6f**). Together, these findings establish that human-specific sequence features at zooHAR.30 sustain FOXC1 occupancy across clinically and evolutionarily relevant skeletal enhancers, directly linking human regulatory divergence to downstream effectors such as *GDF5*.

### Sequence divergence at zooHAR.30 reshapes transcription factor binding

To identify sequence changes underlying the regulatory differences at zooHAR.30, we examined an approximately 380-bp region encompassing annotations from four independent HAR catalogues (**Fig. 7a**). Despite deep evolutionary conservation extending to avian lineages, cross-species alignment identified six human-specific single-nucleotide substitutions together with one additional human-chimpanzee sequence difference within this interval (**Fig. 7c**). We trained ChromBPNet^50^ models on chromatin accessibility data from human embryonic interzone cells and CHON-002 cells. After accounting for Tn5 sequence bias, both models resolved sequence features contributing to predicted accessibility, including conserved motifs for AP-1, a canonical enhancer-binding transcription factor complex (**Fig. 7a-c** and **Supplementary Fig. 9a**).

**Fig. 7.**
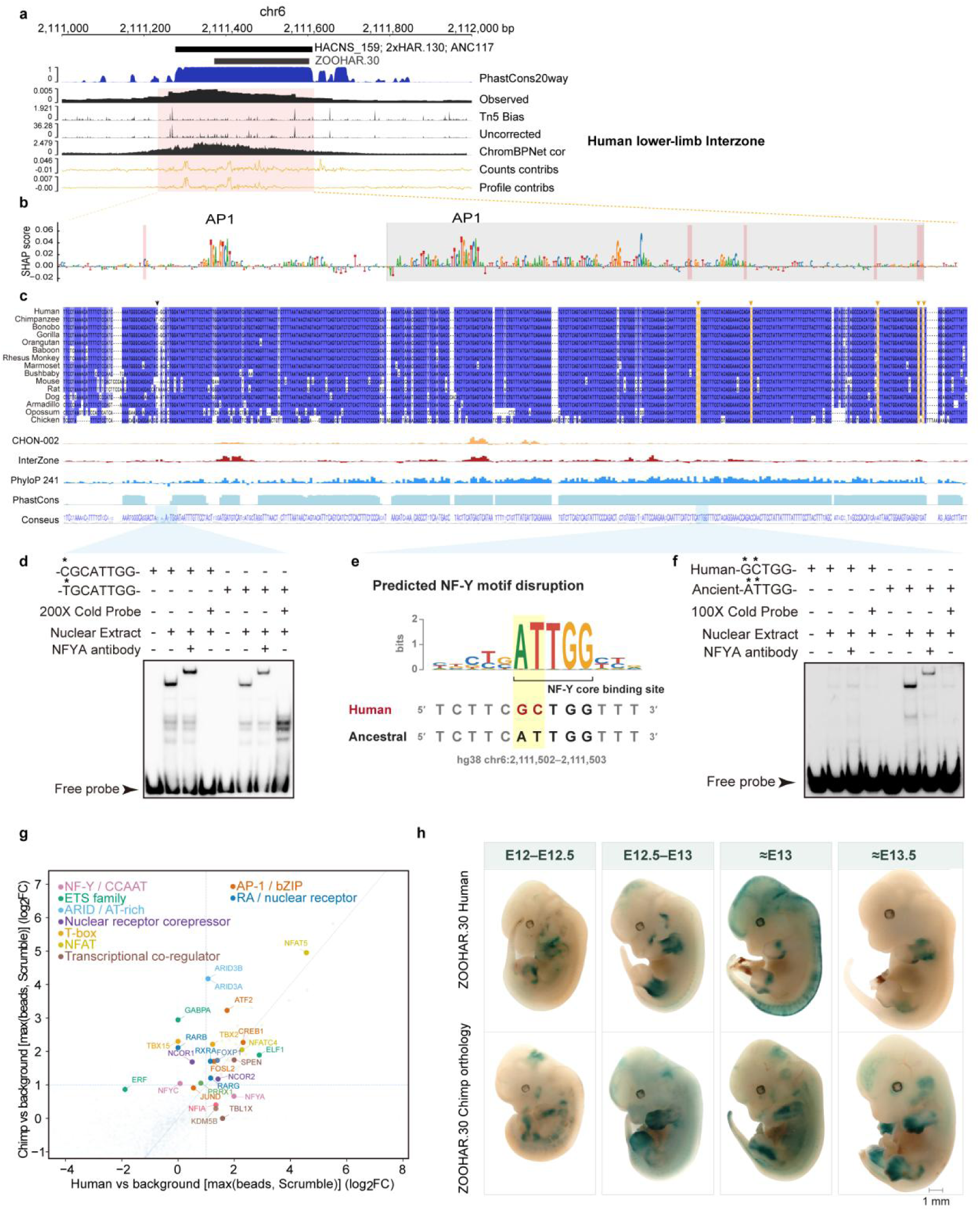
The human zooHAR.30 enhancer reshapes transcription factor binding and exhibits spatially restricted activity in the developing limb. **a**, Chromatin accessibility and ChromBPNet predictions at zooHAR.30 in human lower-limb InterZone cells. Tracks show sequence conservation, observed accessibility, predicted Tn5 bias, uncorrected and bias-corrected accessibility profiles, and base-resolution contribution scores for predicted counts and profiles. Previously annotated human accelerated regions are indicated above the tracks. **b**, Base-resolution contribution scores across the region highlighted in a. Predicted AP-1 motifs are indicated; pink shading marks human–chimpanzee sequence differences. **c**, Alignment of zooHAR.30 sequences from 15 species, with chromatin accessibility tracks from CHON-002 and human InterZone cells and phyloP and phastCons conservation scores. Arrows indicate selected sequence differences. **d**, Electrophoretic mobility shift assay (EMSA) using CHON-002 nuclear extracts and probes containing the indicated human or chimpanzee sequence. A 200-fold excess of unlabeled competitor probe and an NFYA antibody were added as indicated. **e**, Predicted disruption of an NF-Y motif by two adjacent substitutions in the human sequence at hg38 chr6:2,111,502–2,111,503. **f**, EMSA comparing probes containing the human or ancestral sequence at the NF-Y motif. An NFYA-associated complex was detected with the ancestral probe but not the human probe. A 100-fold excess of unlabeled competitor probe and an NFYA antibody were added as indicated. In d and f, the displayed sequences show the variant-containing regions of the probes; arrowheads indicate free probes. **g**, DNA pull-down mass spectrometry using human and chimpanzee zooHAR.30 sequences and CHON-002 nuclear extracts. Each point represents a detected protein or protein group. Axes show log₂ fold enrichment relative to background, defined as the higher signal from the beads-only and scrambled-sequence controls. Colors indicate selected protein groups. **h**, Whole-mount LacZ staining of mouse embryos carrying the human or chimpanzee zooHAR.30 sequence in a piggyBac–Hsp68–lacZ reporter. Both sequences drove reporter activity in developing limbs, with differences in staining patterns between representative embryos. Approximate developmental stages estimated from crown–rump length are indicated. Scale bar, 1 mm. The EMSAs in d and f and the reporter assay in h were each performed in three independent experiments.

We next evaluated each of the seven human–chimpanzee nucleotide differences individually. The divergent site at chr6:2,111,272 (hg38) showed the highest mean signed integrative prioritization score across both models, favoring increased accessibility of the human allele (**Supplementary Fig. 9b** and **Supplementary Table 9**). Electrophoretic mobility shift assays (EMSAs) demonstrated greater NFYA-containing complex formation with the human probe carrying CGCATTGG than with the chimpanzee probe carrying TGCATTGG (**Fig. 7d** and **Supplementary Table 10**). At a second site, two adjacent human-specific substitutions at chr6:2,111,502–2,111,503 converted an ancestral ATTGG sequence matching an NF-Y motif into GCTGG (**Fig. 7e**). EMSA detected an NFYA-containing complex on the ancestral ATTGG probe but not on the human GCTGG probe, indicating loss of NF-Y binding at this position (**Fig. 7f**). Sequence divergence across zooHAR.30 therefore exerts opposing effects on NF-Y binding at two distinct sites. Because NF-Y acts as a pioneer transcription factor that promotes chromatin accessibility^51^, these reciprocal changes in binding may contribute to differences in enhancer activity between the human and chimpanzee alleles.

To interrogate protein interactions across the broader regulatory element, we performed DNA pull-down assays coupled with mass spectrometry using human and chimpanzee zooHAR.30 probes and chondrocyte nuclear extracts. Captured factors included NFYA, the AP-1 subunits FOSL2, JUND and ATF2, the nuclear receptors RARB, RXRA and RARG, and the corepressors NCOR1 and NCOR2 (**Fig. 7g** and **Supplementary Table 10**). A parallel pull-down using mouse limb-bud nuclear extracts recovered NF-Y, AP-1 family members, the nuclear receptors RARB, RXRA and RARG, and the corepressors NCOR1 and NCOR2, together with limb-patterning transcription factors including PRRX1, TBX15, MSX2 and LEF1 (**Supplementary Fig. 9d**). We also assessed CREB5, a transcription factor required for interzone specification across most synovial joints^52^. Reanalysis of published CUT&RUN profiles from E14.5 mouse embryos^53^ revealed CREB5 occupancy at the mouse zooHAR.30 orthologue in knee joint progenitors and in CREB5⁺HOXA11⁺ cells of the developing zeugopod (**Supplementary Fig. 9c**). Together, these findings reveal how evolutionary sequence changes remodel transcription factor binding within an ancestrally conserved enhancer, potentially altering the integration of regulatory inputs governing limb patterning, joint formation and skeletal development.

### The human zooHAR.30 enhancer exhibits spatially restricted activity in the developing limb

We compared the expression patterns driven by the human and chimpanzee zooHAR.30 sequences in transgenic mouse embryos. Each sequence was incorporated into a piggyBac–Hsp68-lacZ reporter construct flanked by insulators to minimize position-dependent effects. Following pronuclear injection into fertilized mouse oocytes, we assessed reporter activity from approximately E11.5 to E13.5, staging embryos by crown-rump length (**Fig. 7h** and **Supplementary Fig. 10**).

Both orthologous sequences drove *lacZ* expression in the posterior limb bud, overlapping the region corresponding to the *Shh*-expressing zone of polarizing activity (**Supplementary Fig. 10**). In stage-matched embryos, activity of the human sequence was largely restricted to the zeugopod, broadly matching the endogenous *Hoxa11* expression domain mapped by in situ hybridization^54,55^, whereas the chimpanzee sequence drove broader expression across the limb bud (**Fig. 7h**). Notably, the zeugopod gives rise to the forearm and lower leg, whose proportions have diverged markedly between humans and other living apes^2,56^.

Beyond the limbs, reporter activity was also detected in the supraorbital region, where the spatial extent and developmental timing of staining differed between the two alleles. At later stages, both sequences drove expression in whisker follicles (**Supplementary Fig. 10**), consistent with the established role of *FOXC1* in hair follicle stem cell maintenance and quiescence^57,58^. Together, these in vivo assays demonstrate that evolutionary sequence changes at zooHAR.30 remodel the spatiotemporal specificity of enhancer activity during limb and craniofacial development, providing a developmental basis for how human-specific regulatory divergence may contribute to morphological differences in these structures.

## Discussion

Human skeletal innovations are thought to have arisen largely through modifications to conserved developmental programs rather than the emergence of entirely new ones. The reorientation of the iliac growth plate and delayed ossification during human pelvic development exemplify how changes in the spatial and temporal organization of developmental processes can shape skeletal adaptations for bipedalism^6^. Our cross-species analyses extend this developmental framework to the limb skeleton. Despite broad conservation of osteochondral cell identities and differentiation trajectories across humans, macaques and mice, we identify human-biased transcriptional programs involving developmental regulators and chromatin-modifying factors. These findings suggest that evolutionary divergence in skeletal development can arise through changes in the regulation of conserved cellular programs without fundamental alterations in lineage organization.

Our comparative chromatin interaction maps provide insights into the regulatory architecture underlying such developmental divergence. Whereas conserved chromatin interactions preferentially involve genes essential for skeletal development, human-gained interactions are associated with recent sequence divergence and increased enhancer connectivity. These observations suggest that evolutionary constraint on core developmental regulatory architecture can coexist with considerable flexibility in enhancer-mediated regulation. The zooHAR.30-containing *FOXC1*-*GMDS* locus illustrates this principle that human-specific sequence changes modify enhancer activity within a conserved enhancer-promoter interaction framework. Notably, the human zooHAR.30 sequence exhibits enhanced regulatory activity in chondrocytes but a more spatially restricted activity domain in the developing limb than the chimpanzee sequence. The preferential activity of the human enhancer in the zeugopod, which gives rise to the forearm and lower leg, raises the possibility that regulatory divergence at this element contributed to the evolution of human limb proportions. More broadly, evolutionary changes in enhancer sequences may modify both the magnitude and spatial deployment of conserved developmental programs, providing a regulatory basis for regional diversification of skeletal morphology.

An important implication of our findings is that evolutionary changes in a single cis-regulatory element can have far-reaching consequences for developmental gene regulation. Endogenous replacement of human zooHAR.30 with the orthologous chimpanzee sequence alters *FOXC1* and *GMDS* expression, reduces osteochondral differentiation potential and diminishes FOXC1 occupancy at numerous skeletal regulatory elements. Among these is the R4 enhancer at the *UQCC1*-*GDF5* locus, a regulatory element previously implicated in knee morphology and osteoarthritis susceptibility^44,49^. The concomitant reduction in FOXC1 occupancy and *GDF5* expression suggests a potential mechanism through which regulatory divergence at zooHAR.30 influences downstream skeletal developmental programs. Such regulatory coupling provides a plausible route by which localized sequence changes can generate coordinated developmental effects across multiple loci. It also raises the possibility that regulatory modifications associated with human skeletal evolution may have consequences for susceptibility to joint disorders in modern populations.

The evolutionary history of zooHAR.30 further highlights the distinction between the origin of sequence changes and their developmental consequences. The accelerated evolution of zooHAR.30 has been attributed to GC-biased gene conversion^19^, and our findings show that substitutions within this ancestrally conserved element alter transcription factor binding and enhancer activity. Thus, sequence changes arising through processes other than adaptive selection may nevertheless generate regulatory variation with developmental consequences. The independent acceleration of the orthologous region in the bipedal jerboa (**Supplementary Fig. 9e**) raises the intriguing possibility of recurrent regulatory evolution at this locus in independently evolved bipedal lineages. Whether these sequence changes produce convergent regulatory or skeletal phenotypes remains to be determined.

Several questions remain concerning the developmental and morphological consequences of zooHAR.30 evolution. Although endogenous sequence replacement establishes a functional contribution of the human sequence to osteochondral cell regulation, the relative contributions of *FOXC1* and *GMDS* to the observed cellular phenotypes remain unresolved. Human-chimpanzee hybrid cell models^13^, combined with allele-specific sequence replacement, could further distinguish cis-regulatory effects from differences in the trans-regulatory environment during osteochondral differentiation. Moreover, endogenous sequence replacement in animal models will be important for determining whether zooHAR.30 divergence alters growth plate dynamics, ossification timing or the relative growth of individual limb bones. Together, our findings highlight how localized regulatory sequence evolution within a conserved developmental architecture can reshape broader gene regulatory programs, providing a mechanistic framework for investigating the genetic basis of human skeletal innovations.

## Methods

### Macaque limb sampling and single-cell RNA sequencing

Hindlimbs were collected from rhesus macaque embryos generated by in vitro fertilization at E35 (*n* = 1), E40 (*n* = 2) and E60 (*n* = 2) under a protocol approved by the Institutional Animal Care and Use Committee of the Kunming Institute of Zoology, Chinese Academy of Sciences (IACUC-PE-2023-11-002), with general anesthesia specified for surgery. Whole hindlimbs were used to construct one library per embryo at E35 and E40, whereas the stylopod and zeugopod were processed separately at E60. Freshly dissected tissues were dissociated into single-cell suspensions, and libraries were prepared by Shanghai Biotechnology Corporation (Shanghai, China) on the Chromium Single Cell 3′ platform (10x Genomics) and sequenced on an Illumina NovaSeq 6000. Library quality metrics and cell-type markers are listed in Supplementary Tables 1 and 2.

### Cross-species integration and cell-type mapping

Macaque limb transcriptomes were integrated with published human and mouse datasets across shared one-to-one orthologs (**Supplementary Table 3**) by fitting SysVI to log-normalized expression of genes highly variable in all three species, with species included as the system covariate. Leiden clustering and UMAP embedding were performed on the resulting latent space, and cell types were annotated from canonical markers together with the original labels. Cross-species correspondence for each human cell type was estimated separately in macaque and mouse from the labels of each cell’s 50 nearest neighbors. Chondrocyte cell-line transcriptomes were projected onto principal components fitted to human developmental pseudobulk profiles using shared orthologs.

### Conserved cell-type expression and skeletal-trait enrichment

Counts were aggregated by cell type and sample (human and mouse) or embryo (macaque). Within each species, limma–voom models compared each cell type against all other lineages, blocking by sample or embryo with equal lineage weights. Conserved markers were defined by log_2_ fold change ≥ 0.5 in all three species and Benjamini–Hochberg false discovery rate (FDR) ≤ 0.05 across gene–cell-type tests, applied to conjunction *P* values (the largest one-sided *P* value across species). Enrichment of skeletal-trait genes from the skeletal-proportion GWAS^3^ (Supplementary Table S20) was evaluated using one-sided Fisher’s exact tests against orthologs testable in all species, with Benjamini–Hochberg correction across cell-type–trait pairs.

### Osteochondral differentiation trajectories

Diffusion pseudotime was computed in Scanpy, rooted at the mesenchymal condensation cell closest to the mean of the three species’ median condensation-cell coordinates. Three branches sharing mesenchymal condensation cells and osteochondral progenitors (OCPs) were defined from diffusion coordinates and cell annotations: the osteoblast, interzone–chondroprogenitor–resting chondrocyte, and perichondrial–articular trajectories. For heatmaps, expression was averaged within pseudotime bins across samples within each species and then across species. For selected genes, species-specific cubic curves were fitted along the combined chondrogenic and joint trajectory, with confidence bands estimated by resampling samples within species.

### Stage matching and cross-species differential expression

Human-equivalent ages for macaque embryos were estimated using kernel ridge regression trained on cell-type proportions and cell-type-specific mean expression from human donors, with inner cross-validation for model selection and leave-one-donor-out evaluation. The early-stage comparison comprised three human donors (post-conception weeks 5.4, 5.6 and 6.1), two E40 macaque embryos and four E12.5 mouse libraries (**Supplementary Table 3**).

Within each cell type, lowly expressed genes were filtered in edgeR and TMM-normalized pseudobulk counts were analyzed with *limma-voom*. For each direction, the larger one-sided *P* value from the human–macaque and human–mouse contrasts was taken, and the smaller directional value was doubled (capped at 1). Differentially expressed genes required concordant effects with |log_2_ fold change| ≥ 0.5 in both contrasts and Benjamini-Hochberg FDR ≤ 0.05 within cell type, and had to be detected in every pseudobulk sample (**Supplementary Table 4**) or in each species for cross-species expression comparisons. Human-higher and human-lower OCP genes were tested separately for KEGG and Reactome enrichment using one-sided hypergeometric tests against all tested OCP genes, with Benjamini–Hochberg correction within each direction.

### Cell lines and culture conditions

Human CHON-002 cells (ATCC, CRL-2847), derived from fetal long-bone cartilage, were maintained in DMEM with 10% heat-inactivated FBS and 0.1 mg/ml G418. Immortalized mouse articular chondrocytes (iMAC; Applied Biological Materials, T0010) were cultured in PriGrow III medium (TM003) with 10% non-heat-inactivated FBS and 1% penicillin–streptomycin. All cultures were kept at 37 °C in 5% CO_2_. Cell lines were authenticated by short tandem repeat (STR) profiling and routinely screened for mycoplasma; parental CHON-002 cells tested negative prior to genome editing.

### Establishment of an immortalized fetal macaque chondrocyte cell line

An immortalized fetal rhesus macaque chondrocyte line was established as previously described^59^. Briefly, primary chondrocytes were isolated from hindlimb long-bone cartilage of an E60 rhesus macaque fetus. After removal of surrounding non-cartilaginous tissue, cartilage was minced and digested sequentially with 0.25% trypsin (30–45 min) and type II collagenase (4–6 h) at 37 °C. Cells were cultured in chondrocyte medium (ScienCell, 4651) containing 5% FBS and 0.5% chondrocyte growth supplement at 37 °C with 5% CO_2_.

Human telomerase reverse transcriptase (hTERT) was cloned into the GV341 lentiviral vector, packaged in HEK293T cells and used to transduce primary chondrocytes at a multiplicity of infection of 1 at 20–30% confluence. Medium was refreshed 16 h post-transduction, and selection with 1.2 μg/ml puromycin began at 72 h. Colonies derived from 2–5 cells were picked, expanded and maintained in 0.3–0.6 μg/ml puromycin.

Expanded lines were characterized by hTERT expression, morphology, G-banded karyotyping, immunostaining and RT–qPCR for chondrocyte markers. The resulting line carried 42 chromosomes and tested negative for mycoplasma before downstream assays. Aggrecan and type II collagen were stained with mouse monoclonal anti-aggrecan (Santa Cruz Biotechnology) and rabbit polyclonal anti-type II collagen (Abcam), followed by Alexa Fluor 488 goat anti-mouse IgG (Abcam) and Alexa Fluor 594 donkey anti-rabbit IgG (Invitrogen). RT–qPCR was run in technical triplicate using AceQ Universal SYBR qPCR Master Mix (Vazyme); primer sequences are provided in Supplementary Table 10.

### HiChIP library preparation and sequencing

H3K27ac and H3K4me3 HiChIP libraries were prepared from human, macaque and mouse chondrocyte cell lines following Mumbach et al.^60^ with minor modifications, using 1 × 10^7^ cells per reaction and two independent biological replicates per species and histone mark.

Cells were crosslinked with 1% methanol-free formaldehyde for 10 min at room temperature and quenched with 125 mM glycine for 5 min. Nuclei were digested with MboI (37 °C, 2 h), filled in with biotin-dATP using Klenow fragment (37 °C, 1 h) and proximity-ligated with T4 DNA ligase (room temperature, 4 h). Chromatin was sheared on a Diagenode sonicator (8–10 cycles, 30 s on / 30 s off) to 200–500 bp, and clarified lysates were diluted up to 1:9 with ChIP dilution buffer to∼0.1% SDS.

Lysates were precleared with Protein A magnetic beads (1 h) and incubated overnight at 4 °C with 7.5 μg anti-H3K4me3 (Abcam, ab8580) or anti-H3K27ac (Abcam, ab4729). Complexes were captured on Protein A beads (2 h, 4 °C) and washed three times each with low-salt, high-salt and LiCl buffers. Following elution, proteinase K digestion and reverse crosslinking, DNA was column-purified and quantified by Qubit.

Biotin-labeled junctions were captured on Streptavidin C1 beads and tagmented on beads at 55 °C for 10 min using 2.5 μl Tn5 per 50 ng DNA (up to 4 μl). Tagmentation was quenched with 50 mM EDTA, and libraries were amplified with Phusion HF master mix and Nextera primers (cycle numbers scaled to post-ChIP DNA input), size-selected to ∼300–700 bp with AMPure XP beads, checked on a Bioanalyzer and sequenced as 150-bp paired-end reads on an Illumina platform.

### Cross-species identification and structural characterization of chromatin interactions

HiChIP reads were processed with HiC-Pro v3.1.0 against hs1 (human), rheMac10 (macaque) or mm10 (mouse). Intrachromosomal interactions were called with HiC-DC+ at 5-kb resolution (FDR ≤ 0.05), excluding bin pairs separated by ≤5 kb. Conserved and shared candidates required overlap at both anchors, whereas species-specific gain candidates lacked a corresponding call in either comparator within a 10-kb tolerance at each anchor.

For each candidate, raw and distance-normalized contacts were evaluated within ±20 bins of the orthologous anchors in each native map. Focal enrichment above axial background and extended axial enrichment were used to classify focal and stripe components, requiring an enrichment score ≥ 1.5 and Holm-adjusted *P* ≤ 0.05 against chromosome-, distance- and coverage-matched controls, as well as recurrence in ≥80% of downsamplings to matched contact depth. Poorly mapped or low-coverage loci were marked unassessable.

Species-specific gains required stronger component support than both other species at full and matched depths. Primate-shared interactions required support in human and macaque with no support in an assessable mouse locus. Conserved interactions required support in all three species, and human-specific losses required support in macaque and mouse paired with depletion, remodeling or absence of the corresponding human component (**Supplementary Table 5**).

### Functional annotation of chromatin interactions

Loop anchors (5 kb) were annotated using species-matched RefSeq genes, public H3K27ac ChIP-seq peaks from ENCODE and ATAC-seq peaks (including in-house CHON-002 ATAC-seq). Promoters (P) were defined as ±2.5 kb around annotated transcription start sites, and enhancers (E) as non-promoter regions overlapping both H3K27ac and ATAC-seq peaks. Promoters took precedence over enhancers; remaining anchors were labeled other (O). Interactions were classified as promoter- or enhancer-connected if either anchor carried that annotation, so P–E interactions contributed to both groups.

### Sequence conservation and human-specific SNV enrichment

Loop anchors were lifted from hs1 to hg38 using UCSC liftOver, and phastCons100way scores were averaged over covered bases within each anchor and then across anchor pairs. Human-gained interactions were compared with primate-shared and conserved interactions by two-sided Wilcoxon rank-sum tests with Benjamini-Hochberg correction.

Candidate human-specific substitutions were obtained from the Ensembl Compara Age of Base annotations (Hsap_ages_1995_105.bb, hg38). Candidates were further screened using UCSC hg38 multiz30way alignments, requiring at least three aligned sequences, including human, and a human base differing from the predominant base among the non-human sequences. To enrich for fixed or nearly fixed human-specific substitutions, candidates were retained if they were absent from gnomAD genomes v3.1.2 or overlapped only variants with alternate allele frequencies ≤0.005 (AC/AN). The filtered SNVs were then lifted over to hs1 coordinates (excluding unmapped or ambiguously mapped loci) for downstream enrichment analysis.

Anchor intervals were merged within each category, and SNVs overlapping the T2T blacklist were excluded. Effective anchor lengths were calculated by subtracting blacklisted bases, and intervals with no remaining bases were discarded. Human-specific SNV density in each category was compared against the pooled remaining categories using negative binomial regression with a log(effective length) offset and adjustment for chromosome and anchor length. Two-sided Wald tests used standard errors clustered by 5-Mb genomic blocks, with Benjamini–Hochberg correction across the six comparisons within each histone mark.

### Alu and hCONDEL enrichment

AluJ, AluS and AluY elements (CHM13v2.0 RepeatMasker) were compared across unique autosomal anchors of human-gained and primate-shared interactions, retaining shared anchors in both sets. Overlap odds ratios, 95% confidence intervals and two-sided *P* values were estimated by chromosome jackknife and Benjamini–Hochberg-adjusted across the three subfamilies. For human-specific deletions, the proportion of loops with an anchor overlapping an hCONDEL junction was compared between human-lost and conserved interactions. Null distributions were generated from 10,000 randomizations matched for chromosome, junction age, conserved-block length, human–chimpanzee divergence, GC content and 20- and 100-way phastCons scores. One-sided empirical *P* values were Benjamini-Hochberg-adjusted across the two HiChIP marks.

### Rare disease group enrichment of conserved interaction target genes

Target genes of conserved interactions were tested for enrichment across Orphanet disease groups against all other chondrocyte-expressed genes. Disease groups were assembled from descendant disorders and germline disease-causing genes in Orphadata (July 2026) and evaluated by one-sided Fisher’s exact tests with Benjamini–Hochberg correction.

### Generation and validation of zooHAR.30-edited CHON-002 cells

Endogenous human zooHAR.30 was replaced with its chimpanzee ortholog in CHON-002 cells by CRISPR–Cas9 homology-directed repair. Cas9–GFP (IDT) was complexed with two sgRNAs flanking zooHAR.30 and co-delivered with an ssODN carrying the chimpanzee sequence flanked by human homology arms (Supplementary Table 10). Electroporation was carried out on a Neon Transfection System (Thermo Fisher Scientific) in 10-μl reactions (5 × 10^5^ cells, 2 μM Cas9–GFP, 1 μM each sgRNA, 1 μM ssODN; 1,600 V, five 10-ms pulses). Cells were recovered for 24 h in antibiotic-free complete medium containing 1 μM Alt-R HDR Enhancer V2 (IDT), sorted for GFP positivity by flow cytometry and expanded in prewarmed medium for 2 weeks before single-cell cloning. Because CHON-002 cells exhibit low single-cell clonogenicity, ScienCell chondrocyte medium was used until visible colonies emerged, after which clones were returned to routine medium. zooHAR.30-deletion (HAR^KO^) cells were generated using the same sgRNA pair without the ssODN template. Non-targeting control lines were generated in parallel using non-targeting sgRNAs.

Genomic DNA from expanded single-cell clones was isolated with the TaKaRa MiniBEST Universal Genomic DNA Extraction Kit Ver.5.0 (9765) following the animal-tissue lysis protocol. Amplicons spanning the edited locus and flanking regions were sequenced by NGS to confirm the intended edit, and the top 10 CRISPOR-predicted off-target sites per sgRNA were PCR-amplified and sequenced to rule out off-target editing. H3K27ac CUT&Tag was performed in non-targeting control, Chimp-HAR^KI^ and HAR^KO^ cells. Multiple independently derived clonal lines were established for each genotype (non-targeting control, Chimp-HAR^KI^ and HAR^KO^). Clones with verified intended edits and no detectable off-target mutations were selected for all subsequent downstream experiments.

### Cell proliferation and apoptosis assays

EdU incorporation was assayed with the BeyoClick EdU-488 kit (Beyotime, C0071L). Chimp-HAR^KI^ and non-targeting control CHON-002 cells were seeded at matched densities (3–5 × 10^5^ cells per 60-mm dish), cultured for 24 h and pulsed with 10 μM EdU for 4 h at 37 °C. EdU-positive fractions were quantified by flow cytometry using an unlabeled control to set gates; three independent cultures per group were compared by two-sided Welch’s *t*-test.

For CCK-8 assays (Proteintech), 1,000 cells were seeded per well in 96-well plates in 100 μl complete medium. At 24, 48, 72, 96 and 120 h, 10 μl reagent was added per well for 1–2 h at 37 °C, and blank-corrected absorbance was read at 450 nm. Groups were compared at each time point using two-sided Welch’s *t*-tests.

Apoptosis was measured with the Annexin V Alexa Fluor 488/propidium iodide kit (Thermo Fisher Scientific, V13241). Cells (1 × 10^5^ per sample) were stained for 15 min at room temperature and analyzed by flow cytometry, quantifying Annexin V^+^/PI^−^ and Annexin V^+^/PI^+^ populations separately across three independent replicates per group (compared by two-sided unpaired *t*-tests).

### Osteogenic and chondrogenic differentiation assays

For two-dimensional osteogenic induction, Chimp-HAR^KI^ and control CHON-002 cells were seeded at equal densities in 12-well plates and cultured for 10 days in high-glucose DMEM (with GlutaMAX and sodium pyruvate) supplemented with 10% FBS, 1% penicillin–streptomycin, 100 nM dexamethasone, 50 μg/ml L-ascorbic acid-2-phosphate, 10 mM β-glycerophosphate, 50 ng/ml recombinant human BMP-2 and 10 nM calcitriol. ALP staining and activity were measured with BCIP/NBT and ALP activity kits (Beyotime, C3206 and P0321S), calculating activity from 405-nm absorbance against a *p*-nitrophenol standard curve. ALP activity was compared across six wells per group from one of three independent experiments; osteogenic *ALPL* RNA-seq expression was compared across three control and four Chimp-HAR^KI^ clones (both two-sided Welch’s *t*-tests).

For three-dimensional chondrogenic pellet culture, 1 × 10^5^ cells per well were pelleted in Corning ultra-low attachment 96-well plates (300 × *g*, 3 min) and maintained for 21 days under 5% O_2_ in high-glucose DMEM with GlutaMAX and sodium pyruvate (Gibco, 10569010), 20 mM HEPES, 40 μg/ml L-proline, 50 μg/ml L-ascorbic acid-2-phosphate, 1% ITS+ Premix, 100 nM dexamethasone, 10 ng/ml recombinant human TGF-β3 and 1% penicillin–streptomycin, with half-medium changes every other day. At day 21, chondrospheres were imaged under identical settings for diameter measurement (eight control versus six Chimp-HAR^KI^ chondrospheres from one of three independent experiments; two-sided Welch’s *t*-test).

### RNA extraction and transcriptome analysis

Total RNA was extracted with TRIzol–chloroform; chondrospheres were first homogenized in TRIzol using a 2-ml KIMBLE KONTES Dounce grinder. After quality control, libraries were prepared and sequenced in paired-end mode. Cross-species cell-line comparisons included three samples per species; edited CHON-002 comparisons comprised three control and four Chimp-HAR^KI^ libraries before differentiation, and three control and six Chimp-HAR^KI^ libraries after chondrosphere culture.

Reads from edited cells and chondrospheres were quantified against T2T-CHM13 v2.0 transcripts using Salmon v1.8.0. Genes with ≥10 counts in at least two samples were tested in DESeq2 v1.38.3, requiring |log_2_ fold change| ≥ 1 with Benjamini–Hochberg FDR < 0.01 (undifferentiated cells) or < 0.05 (chondrospheres). GO biological process enrichment was performed by GSEA on undifferentiated-cell genes ranked by log_2_ fold change (clusterProfiler v4.6.2, org.Hs.eg.db v3.16.0; nominal *P* < 0.05 and FDR < 0.05; **Supplementary Table 7**). Cross-species cell-line contrasts are reported in Supplementary Table 6.

### Training of chondrocyte-specific ChromBPNet models

Two independent ChromBPNet models were trained: one on paired-end CHON-002 ATAC-seq generated in this study and one on aggregated interzone-cell accessibility profiles from the human fetal skeletal multiome atlas(E-MTAB-14385). Models were trained with five-fold chromosome-based cross-validation using blacklist-filtered peaks, GC-matched non-peak regions and separately trained Tn5 bias models, taking 2,114-bp sequence inputs to predict total counts and cleavage profiles over the central 1,000 bp.

Bias-corrected models were interpreted across zooHAR.30 with DeepSHAP for both count and profile outputs. De novo motifs identified from count contribution scores with TF-MoDISco were matched against JASPAR vertebrate motifs. Variant contribution scores are listed in Supplementary Table 9.

### Transgenic mouse enhancer assays and whole-mount X-gal staining

Human zooHAR.30 and its chimpanzee ortholog were cloned upstream of a minimal *Hsp68* promoter and *lacZ* reporter in a PiggyBac vector flanked on each side by two copies of a β-globin insulator to buffer position effects. Reporter constructs were co-injected with a CAG-driven PiggyBac transposase helper plasmid into pronuclei of fertilized C57BL/6J embryos. All transgenic mouse procedures were approved by the Institutional Animal Care and Use Committee of Cyagen Biosciences (Suzhou) Inc. (IACUC nos. TACU26-MS001-1660 and TACU26-MS001-1661).

Embryos were harvested between E11 and E13.5 and stained for X-gal following Blanco et al.^61^ with modifications. Embryos were rinsed in cold PBS, fixed on ice in 0.125% glutaraldehyde (duration scaled to size), washed in PBS, incubated for 10 min in X-gal rinse buffer at room temperature and stained in 1 mg/ml X-gal, 5 mM potassium ferricyanide and 5 mM potassium ferrocyanide at 37 °C in the dark for 2–6 h with periodic monitoring. Specimens were then washed in PBS, postfixed in 4% paraformaldehyde at 4 °C and imaged as whole mounts. A spatial domain was scored positive when reproduced in at least two independent embryos carrying the same construct at the same stage across three independent experiments.

### FOXC1 CUT&Tag library preparation

FOXC1 CUT&Tag libraries were prepared from non-targeting control and Chimp-HAR^KI^ CHON-002 cells (two biological replicates per condition) using a Vazyme CUT&Tag kit. Cells were bound to concanavalin A magnetic beads and incubated sequentially with rabbit polyclonal anti-FOXC1 (Invitrogen, PA1-807, 2 μg per reaction) and a secondary antibody, with matched IgG controls processed in parallel. Following pA/G–Tn5 binding, washes and tagmentation, DNA was purified, PCR-amplified with indexed primers and sequenced in paired-end mode.

### FOXC1 CUT&Tag data processing and differential-binding analysis

Paired-end reads were trimmed with fastp v0.12.4 and aligned to T2T-CHM13 v2.0 using Bowtie2 v2.2.5 (--very-sensitive-local, 10–700-bp fragments, concordant pairs only). BAM files were sorted with SAMtools v1.9, and replicate peaks were called against matched IgG controls in MACS3 (paired-end mode, retaining duplicates, *q* < 0.05).

For visualization, replicate BAM files were merged per condition, converted to 10-bp RPGC-normalized BigWig tracks with deepTools bamCoverage and plotted using pyGenomeTracks v3.9.

Differential binding was tested with DiffBind v3.20.0 and DESeq2 v1.50.2 (R v4.5.3). Consensus regions present in at least two of four libraries were quantified in 501-bp summit-centered windows, normalized to full library sizes and filtered at FDR < 0.05 and |log_2_ fold change| ≥ 1 (**Supplementary Table 8**). Regions with higher signal in controls were termed NT-gained and annotated against CHM13 v2.0 with ChIPseeker v1.46.1 (±3-kb promoter window).

### Overlap enrichment of NT-gained FOXC1 peaks in human chondrocyte-accessible chromatin

NT-gained FOXC1 peaks were intersected with two human chondrocyte ATAC-seq references: combined E59 proximal and distal femur, tibia, humerus and radius peaks (GSE122877; GSM3733315–GSM3733322) and the published primary knee articular chondrocyte consensus peak set^45^ (GSE261280_Chondrocytes.idr.optimal_peak.narrowPeak.gz), which was lifted from GRCh38 to hs1 via UCSC liftOver and merged with BEDTools. Each FOXC1 peak overlapping at least one ATAC-seq interval in hs1 was counted once per reference, and NT-gained peaks were compared against the remaining tested regions by two-sided Fisher’s exact tests in R 4.2.2.

### Enrichment of trait-associated variants in differential FOXC1-binding regions

Variant enrichment was tested across NT-gained FOXC1 regions against the 114,520 remaining tested regions. Peak coordinates were lifted from hs1 to GRCh38 (hs1ToHg38, minimum mapping fraction 0.95). European-ancestry 95% credible sets were obtained from Supplementary Table 9 of the osteoarthritis meta-analysis^46^ (hip osteoarthritis) and Supplementary Table S5 (sheet “UKB Leg”) of the skeletal-proportions GWAS^47^ (leg length, UK Biobank SuSiE), with GRCh37 coordinates converted to GRCh38 (hg19ToHg38, default parameters). Unmapped records and variants outside chromosomes 1-22 and X were excluded.

Each region overlapping at least one credible-set variant was counted once per trait. Foreground and background regions were jointly stratified by chromosome and decile of overall FOXC1 signal; fold enrichment was computed as observed foreground hits divided by the exact expectation under stratified sampling without replacement. One-sided empirical *P* values were derived from 100,000 stratified hypergeometric simulations with an add-one correction, alongside exact *P* values obtained by convolving stratum-specific distributions. Each *P*-value set was Benjamini-Hochberg-adjusted across the two traits. Simulations were run in Python 3 (NumPy) and plotted in R 4.2.2.

### Electrophoretic mobility shift and supershift assays

Nuclear extracts were prepared from CHON-002 cells with NE-PER reagents (Thermo Scientific) and quantified by Pierce BCA assay (23225). Matched 5′-biotin-labeled human and chimpanzee double-stranded probes (28 or 35 bp, centered on each candidate variant; Supplementary Table 10) were assayed using the LightShift Chemiluminescent EMSA Kit (Thermo Scientific, 20148). Each 20-μl reaction contained 7 μg nuclear extract, 20 fmol labeled probe, 1× binding buffer, 5% glycerol, 50 ng/μl poly(dI·dC) and 0.05% NP-40, incubated for 20 min at room temperature. For competition, a 200-fold molar excess of unlabeled probe was preincubated with extract for 10 min before adding labeled probe. For supershifts, 1 μl anti-NFYA antibody (Santa Cruz Biotechnology, sc-17753X) was added for an additional 15 min. Complexes were resolved on 6% native polyacrylamide gels, transferred to positively charged nylon membranes, UV-crosslinked and detected by chemiluminescence. Each assay was repeated in three independent experiments.

### DNA affinity purification and mass spectrometry

DNA pull-downs were performed using nuclear extracts from CHON-002 cells and E12 mouse limb buds. Equal-length human and chimpanzee zooHAR.30 probes and a length- and base-composition-matched scrambled control (**Supplementary Table 10**) were PCR-amplified with 5′ Biotin-TEG primers, column-purified, quantified by Qubit dsDNA assay, denatured at 95 °C for 5 min and reannealed by cooling to 25 °C at 0.1 °C/s.

For each reaction, 5–10 μg DNA was immobilized on 50 μl MyOne Streptavidin C1 beads, with beads-only controls run in parallel. Nuclear extracts (500 μg per reaction) were diluted in salt-free binding buffer (20 mM HEPES pH 7.9, 1.5 mM MgCl_2_, 0.2 mM EDTA, 10% glycerol, 1 mM DTT, protease inhibitors and 10 μg poly (dI·dC)), precleared with uncoupled beads (20–30 min, 4 °C) and incubated with probe-bound beads (2–3 h, 4 °C). Beads were washed once with 100 mM KCl / 0.05% NP-40, twice with 150 mM KCl / 0.1% NP-40 and twice with 50 mM ammonium bicarbonate. Bound proteins were eluted in SDS loading buffer (95 °C, 10 min), resolved by short-run SDS–PAGE and excised for GeLC–MS/MS.

Peptides were analyzed on a nanoElute coupled to a timsTOF Pro mass spectrometer (Bruker) in PASEF mode (30-min gradient, 300 nl/min). Raw files were searched in MaxQuant v2.7.0.0 against species-matched UniProt databases (uniprot_Homo_sapiens_205317_20260105 and uniprot_Mus musculus_87600_20260114), allowing two missed tryptic cleavages, fixed cysteine carbamidomethylation, variable methionine oxidation and N-terminal acetylation, 20 ppm precursor and 0.1 Da fragment mass tolerances, 1% FDR and iBAQ quantification. After removing keratins, enrichment was computed relative to the higher signal of the beads-only and scrambled-probe controls, and a 1.5-fold ratio between human and chimpanzee probes was used to separate sequence-biased from shared binders (**Supplementary Table 10**).

### Luciferase reporter assay

The SW1353 (a human chondrosarcoma cell line) and iMAC (an immortalized mouse articular chondrocyte cell line) cell lines were purchased commercially (Servicebio). Both cell lines were cultured in Dulbecco’s Modified Eagle Medium (DMEM) supplemented with 10% fetal bovine serum (FBS) at 37 °C in a humidified incubator containing 5% CO₂. Mycoplasma contamination was routinely monitored using PCR-based assays.

The reference and alternative allele sequences of the predicted enhancers were cloned into the pGL3-promoter vector (Promega, E1761). SW1353 and iMAC cells were seeded in 24-well plates and co-transfected with 500 ng of pGL3-ref/alt-promoter or pGL3-promoter control plasmid and 25 ng of pRL-TK plasmid (Promega, E2241) using Lipofectamine 3000 (Invitrogen, L3000015). After 48 h, cells were washed one time with phosphate-buffered saline (PBS), and luciferase activity was measured using the Dual-Luciferase Reporter Assay System (Promega, E1910) according to the manufacturer’s instructions. Each group included six technical replicates, and the experiment was independently repeated three times. Firefly luciferase activity was normalized to Renilla luciferase activity. Statistical significance was determined using a two-tailed test. ns, P > 0.05; *P < 0.05, **P < 0.01, ***P < 0.001, ****P < 0.0001.

### Statistics and reproducibility

Single-cell sample and library metadata are summarized in Supplementary Tables 1 and 3. Stage-matched differential expression used pseudobulk profiles per cell type, run in R 4.5.0 (edgeR 4.4.2, limma 3.62.2) for early-stage cross-species comparisons and in R 4.3.3 (edgeR 4.0.16, limma 3.58.1) for the broader cell-type analysis (**Supplementary Table 4**). Experimental units, replicate counts and two-sided statistical tests for cellular assays are given in each corresponding section. Embryonic enhancer activity was assessed for reproducibility within each construct and developmental stage.

## Acknowledgements

This study was supported by grants from the National Key Research and Development Program of China (2025YFC3410300 to B.S.), Major Scientific Project of Yunnan Province of China (202305AH340007 to B.S.), National Natural Science Foundation of China (32288101 to B.S.), Yunnan Scientist Workshop (to B.S.), and Key Basic Research Project of Yunnan Province of China (202401BC070014 to B.S.).

## Author contributions

B.S. and Y.Z. conceived and designed the study. Y.Z. performed all data analyses, conducted experiments involving genome-edited cell lines, performed EMSA and DNA pull-down, and prepared the human and mouse HiChIP libraries. J. Yuan performed the dual-luciferase reporter assays. X.Z. performed the ATAC-seq and FOXC1 CUT&Tag experiments. Y.Z., J. Yuan, and X.Z. performed the *LacZ* reporter assays. C.L. and Y.W. established the immortalized macaque hindlimb chondrocyte lines and prepared the macaque HiChIP libraries. J. Yin assisted with cell culture. F.Z. assisted with sample collection for macaque single-cell RNA-seq. B.S. supervised the study. Y.Z. and B.S. wrote the manuscript.

## Data availability

Public datasets analyzed in this study include limb single-cell atlases for human^8^ (E-MTAB-8813) and mouse (GSE275855, ENCSR713GIS and E-MTAB-10514), the human fetal skeletal multiome atlas^7^ (E-MTAB-14385), human limb spatial transcriptomics^8^ (E-MTAB-10367), chondrocyte ATAC-seq references (GSE122877^44^ and GSE261280^45^), developmental limb H3K27ac ChIP-seq data from human, rhesus macaque and mouse^12^ (GSE42413), and mouse embryonic CREB5 CUT&RUN data^53^ (GSE281189, a subseries of GSE281192). Public chondrocyte ChIP-seq datasets for H3K27ac (ENCSR925IAP), H3K4me3 (ENCSR434WEY) and CTCF (ENCFF835UIE) were obtained from ENCODE. Sequencing datasets generated in this study will be deposited in public repositories upon peer-reviewed publication and are available from the corresponding author upon reasonable request prior to publication. Processed summary results, metadata and oligonucleotide sequences are provided in Supplementary Tables 1-10.

## Code availability

Custom analysis scripts used in this study will be made publicly available upon peer-reviewed publication and are available from the corresponding author upon request.

## Supplementary Figures

**Supplementary Fig. 1.**
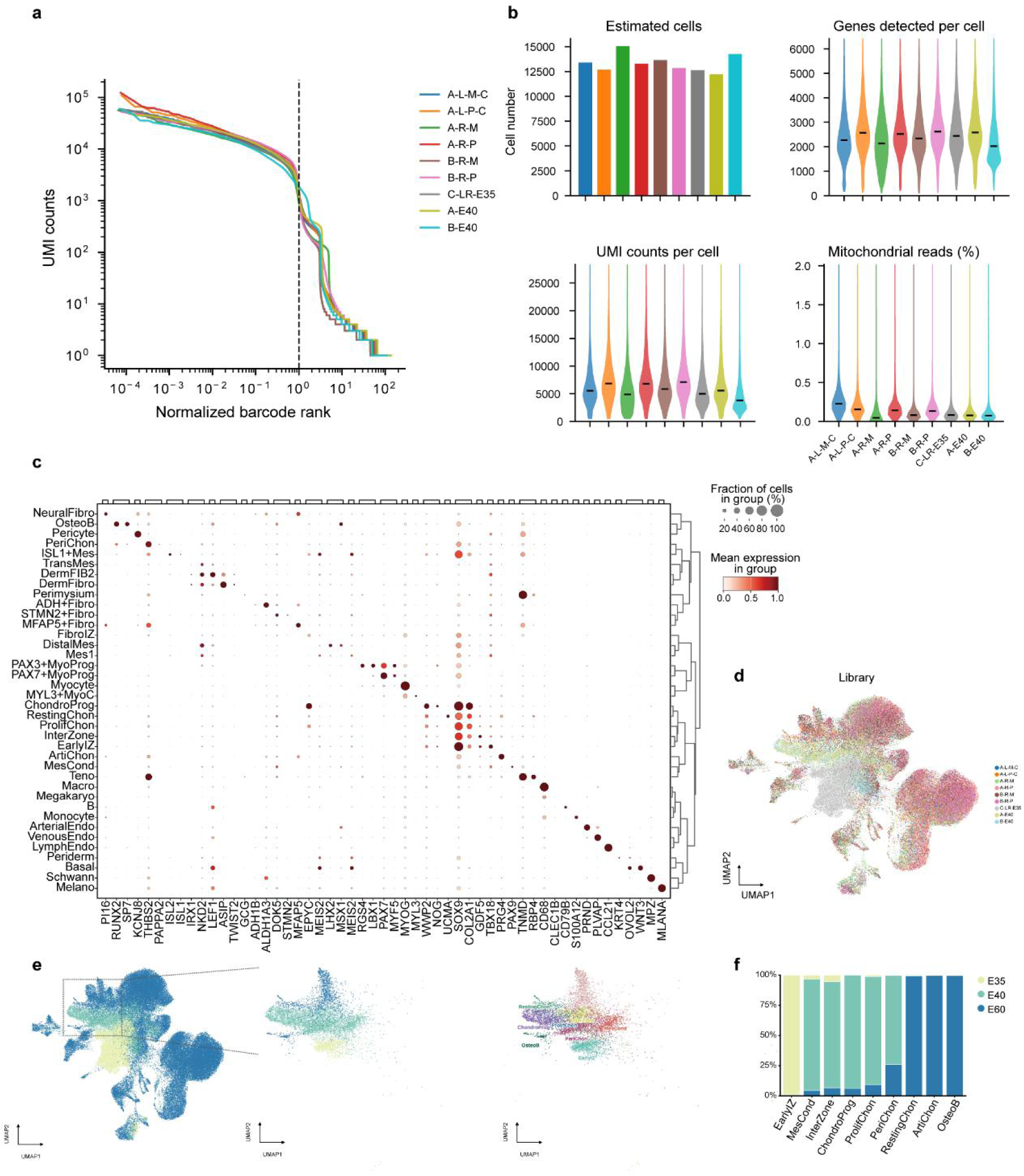
Single-cell transcriptomic profiling of rhesus macaque embryonic hindlimbs. **a,** Barcode-rank curves showing unique molecular identifier (UMI) counts across nine libraries. Barcode ranks were normalized to the number of cells identified in each library; the dashed line indicates a normalized rank of 1. **b,** Estimated cell numbers per library and distributions of detected genes, UMI counts and mitochondrial read percentages per cell. Horizontal lines indicate medians. **c,** Expression of selected marker genes across annotated cell types. Dot size indicates the fraction of cells expressing each gene, and color indicates mean expression. **d,** UMAP of macaque hindlimb cells colored by library. **e,** UMAP highlighting the osteochondral lineage, with selected cells shown by developmental stage and annotated cell type. **f,** Proportions of cells from each developmental stage within the indicated osteochondral cell types.

**Supplementary Fig. 2.**
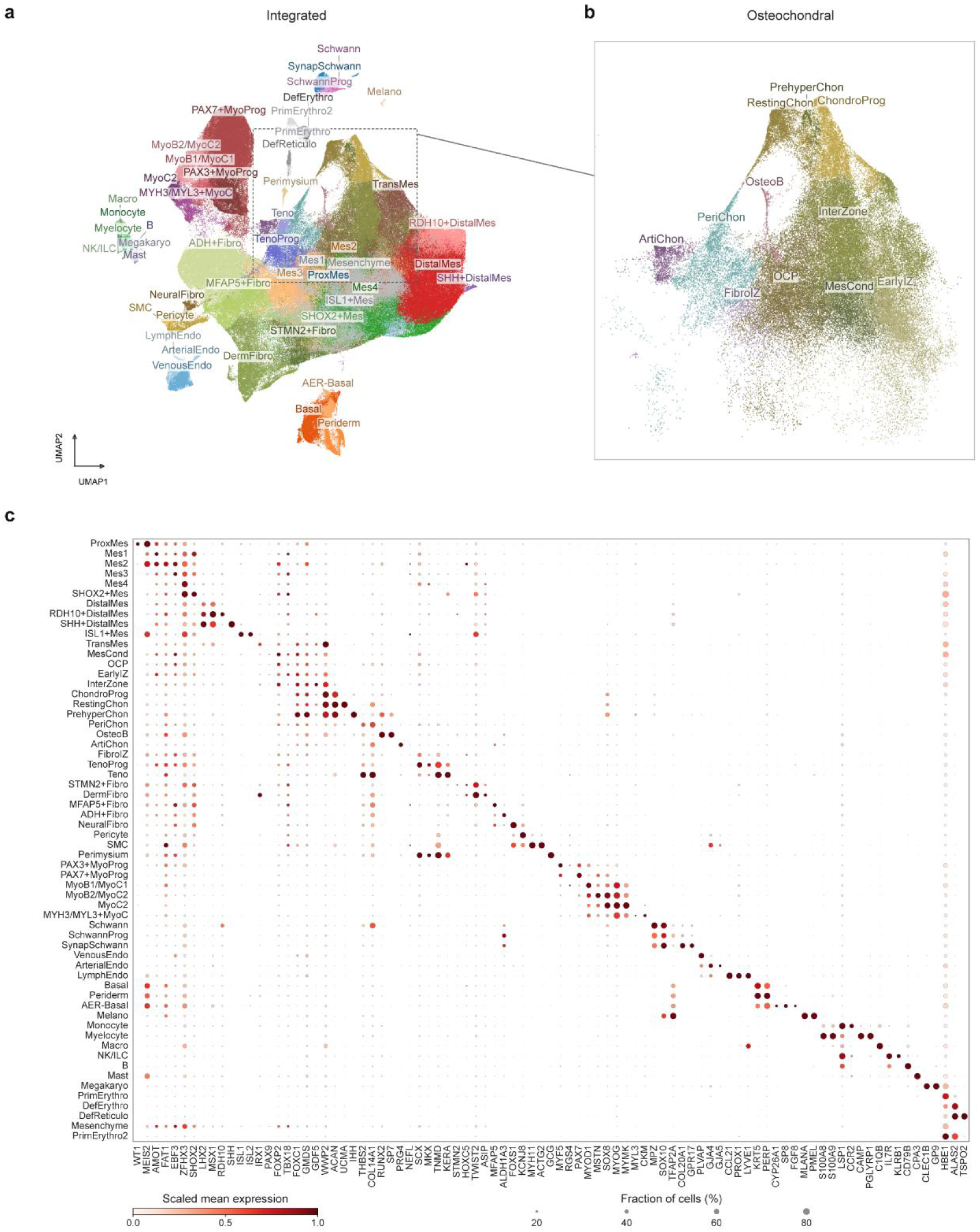
Cell-type annotation of the integrated cross-species limb atlas. **a,** UMAP of single-cell transcriptomes from developing human, macaque and mouse limbs integrated using sysVI and one-to-one orthologous genes. Cells are colored by annotated cell type; each dot represents one cell. **b,** Enlarged view of the osteochondral cell populations in the boxed region in **a**, shown in the same UMAP embedding and colored by cell type. **c,** Dot plot showing the expression of selected marker genes across annotated cell types in the integrated atlas. Dot size indicates the percentage of cells expressing each gene within a cell type, and color indicates scaled mean expression.

**Supplementary Fig. 3.**
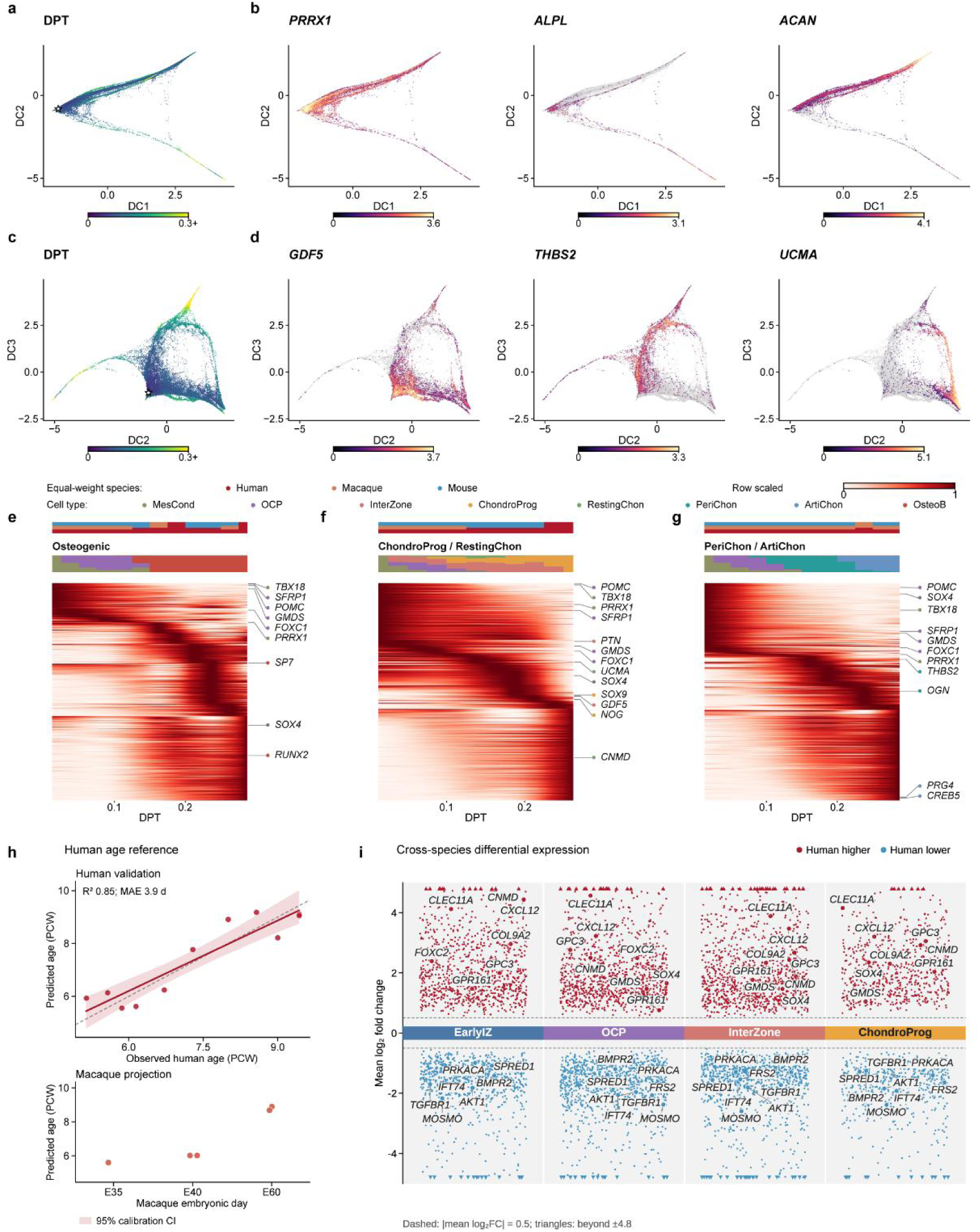
Osteochondral differentiation trajectories and cross-species differences in gene expression. **a,** Diffusion map of integrated osteochondral lineage cells projected onto diffusion components (DCs) 1 and 2 and colored by diffusion pseudotime (DPT). The star indicates the root MesCond cell. **b,** Expression of *PRRX1*, *ALPL* and *ACAN* projected onto the embedding in **a**. **c,** Projection of the same cells onto DC2 and DC3, colored by DPT. The star indicates the root cell. **d,** Expression of *GDF5*, *THBS2* and *UCMA* projected onto the embedding in **c**. In **a–d**, DC coordinates were inverse hyperbolic sine-transformed, and the sign of DC2 was reversed for display.**e-g,** Heatmaps showing gene-expression dynamics along the osteogenic (**e**), chondroprogenitor and resting chondrocyte (**f**), and perichondrial and articular chondrocyte (**g**) trajectories. Expression within each DPT bin was averaged with equal weights for samples within each species and for the species represented, smoothed and scaled from 0 to 1 for each gene. The upper bars indicate species weights and cell-type composition, respectively. **h,** Validation of a human age-reference model using nested leave-one-donor-out predictions for 11 human donors (top) and projection of five macaque embryos onto the human reference (bottom). The model incorporates cell-type composition and gene expression. The solid line indicates the linear calibration fit; shading indicates its 95% pointwise bootstrap confidence interval. The dashed line indicates the identity line. *R*² and mean absolute error (MAE) were calculated from held-out human predictions. PCW, post-conception weeks. **i,** Pseudobulk differential-expression analysis comparing human with macaque and mouse in early interzone cells (EarlyIZ), osteochondral progenitors (OCP), interzone cells (InterZone) and chondroprogenitors (ChondroProg). Each point represents a gene; its vertical position indicates the mean log₂ fold change across the human–macaque and human-mouse comparisons. Displayed genes showed concordant changes in both comparisons, with |log₂ fold change| ≥ 0.5 in each comparison and joint false discovery rate ≤ 0.05. Red and blue indicate higher and lower expression in human, respectively. Selected developmental genes with concordant changes in at least two cell types are labelled. Dashed lines indicate mean log₂ fold changes of ±0.5; triangles indicate values beyond ±4.8.

**Supplementary Fig. 4.**
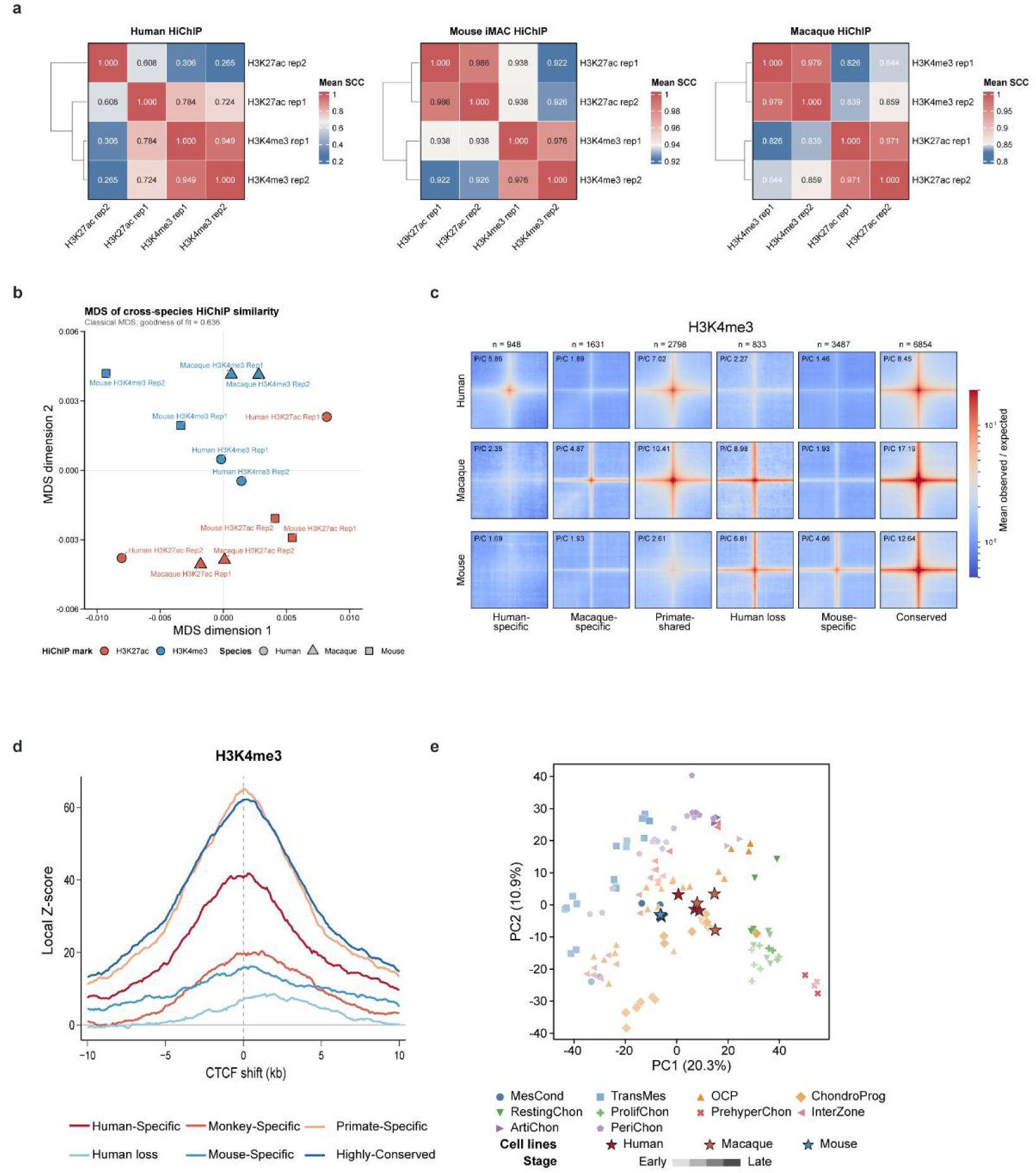
Characterization of cross-species HiChIP datasets and chondrocyte cell lines. **a,** Heatmaps of mean stratum-adjusted correlation coefficients (SCCs) between H3K27ac and H3K4me3 HiChIP replicates within each species. **b,** Multidimensional scaling (MDS) of HiChIP samples from the three species. Colors indicate histone marks, and symbols indicate species. **c,** Aggregate H3K4me3 HiChIP contact maps for six interaction categories in human, macaque and mouse. Colors show mean observed-to-expected contact frequency. Numbers above the maps indicate interactions; P/C indicates the ratio of the central signal to the corner background. **d,** Local Z-scores for the overlap between CTCF sites and H3K4me3 HiChIP anchors across interaction categories as CTCF sites were shifted relative to the anchors. **e,** Principal component analysis of human developmental cell-type pseudobulk profiles, with human, macaque and mouse chondrocyte cell-line transcriptomes projected onto the developmental reference. Symbols and colors indicate cell types, shading indicates developmental stage, and stars indicate cell-line samples.

**Supplementary Fig. 5.**
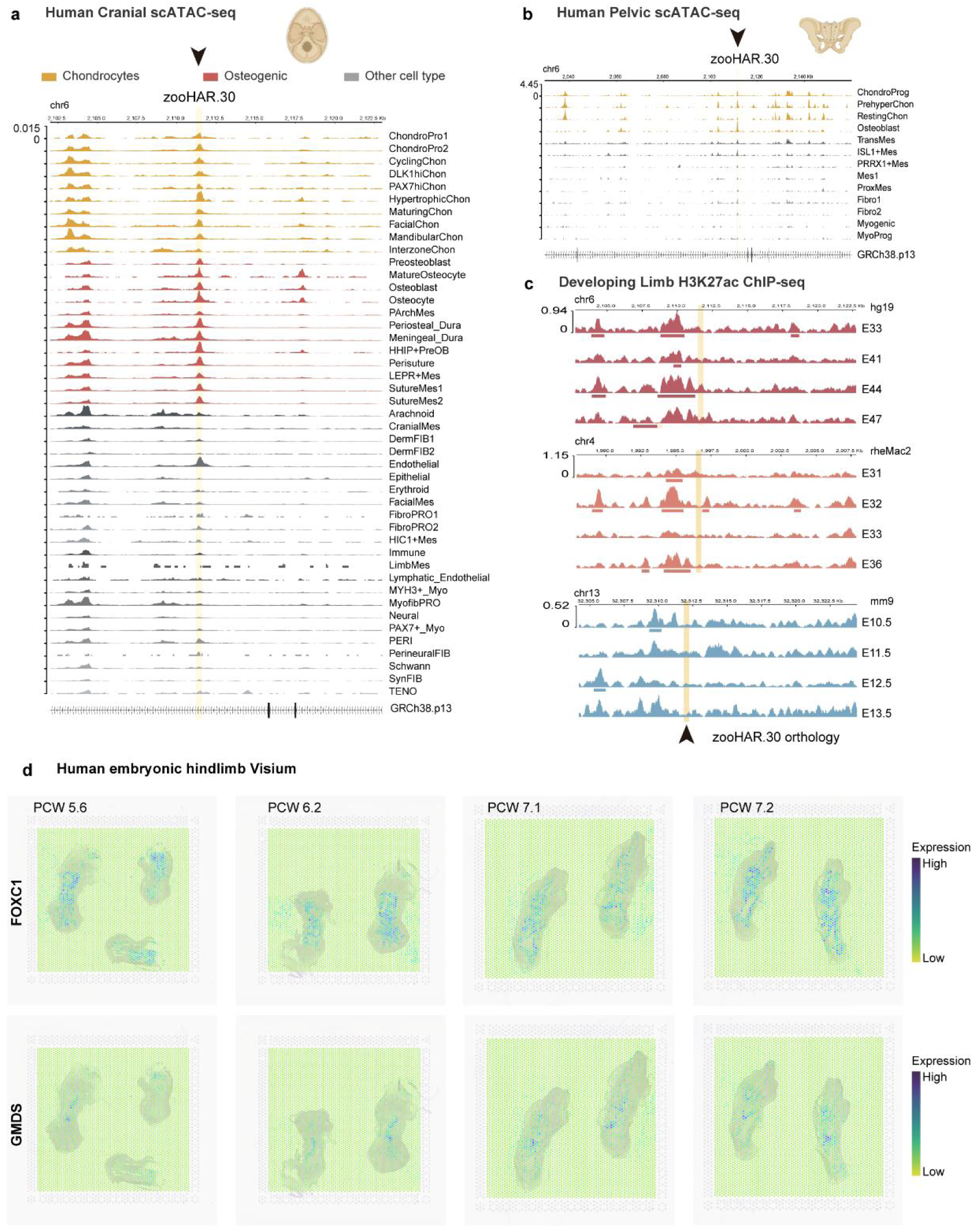
Chromatin accessibility and developmental activity at zooHAR.30. **a**,**b**, Cell-type-resolved chromatin accessibility at zooHAR.30 in human developing cranial skeletal (a) and pelvic (b) tissues. Colored tracks highlight chondrogenic and osteogenic cell types; grey tracks represent other cell types. Yellow highlights indicate zooHAR.30. **c**, H3K27ac profiles at zooHAR.30 and its orthologous regions in developing human, macaque and mouse limbs. Shaded regions indicate the corresponding sequence in each species. **d**, Spatial expression of FOXC1 and GMDS in human embryonic hindlimb sections at the indicated post-conception weeks (PCW), assessed by Visium spatial transcriptomics.

**Supplementary Fig. 6.**
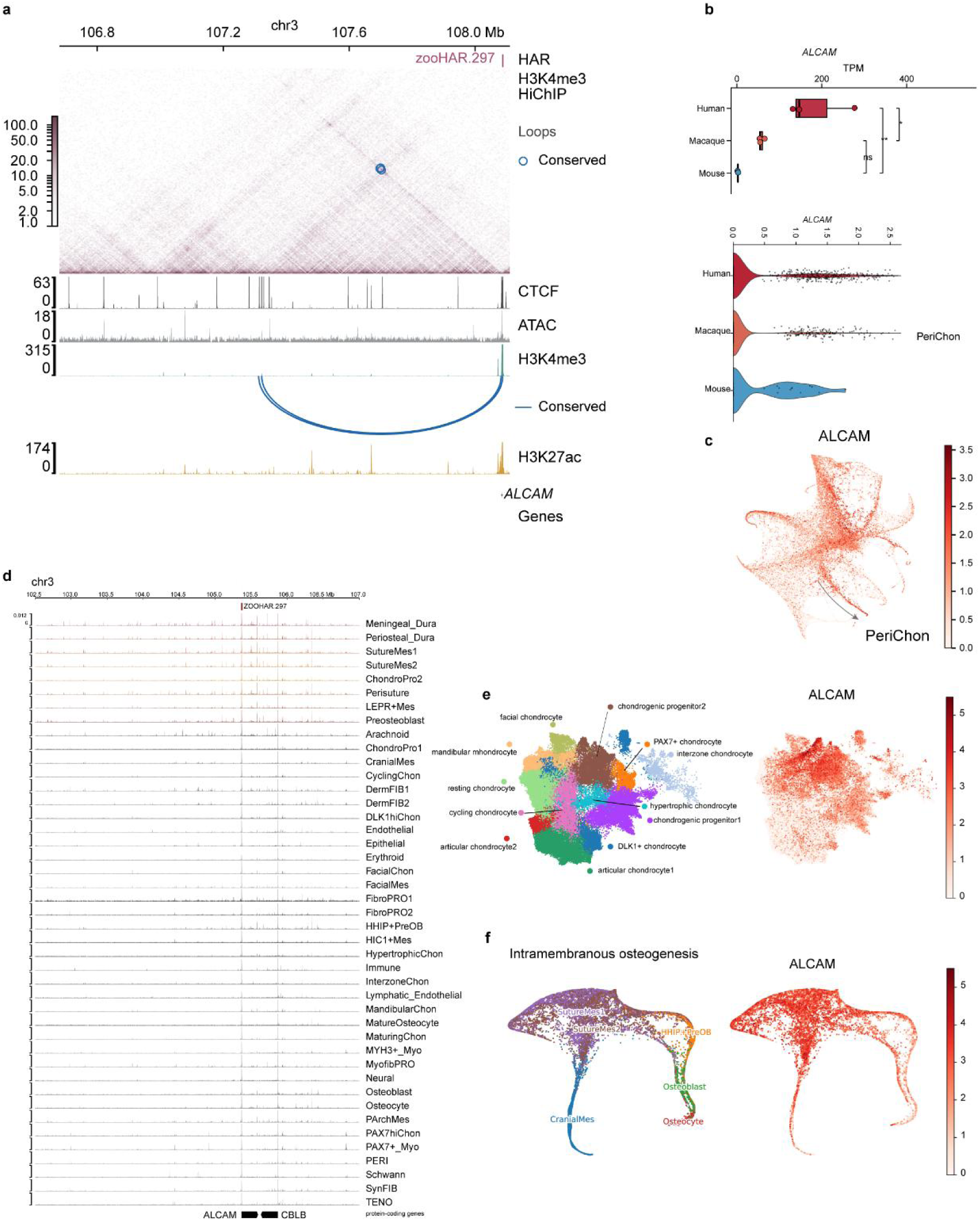
zooHAR.297 resides within a conserved regulatory landscape at the *ALCAM* locus active in perichondrial and cranial progenitors. **a**, Human H3K4me3 HiChIP contact map alongside CTCF, ATAC-seq, H3K4me3 and H3K27ac profiles across the *ALCAM* locus, which encodes the cell adhesion molecule CD166 involved in osteochondral progenitor regulation^34^. Species-conserved chromatin interactions (blue arcs) anchor near the *ALCAM* promoter and overlap zooHAR.297 as well as a CTCF-occupied site. Genomic coordinates are shown in the CHM13 assembly**. b**, Expression of *ALCAM* across human, macaque and mouse chondrocyte lines (top) and in stage-matched developing limb perichondrial cells (PeriChon; bottom), demonstrating significantly elevated *ALCAM* expression in human chondrocytes relative to non-human primates and mice, as well as conserved expression in PeriChon progenitors in vivo.In the top panel, boxes indicate the median and interquartile range, whiskers extend to the most extreme values within 1.5× the interquartile range, and points represent individual RNA-seq samples (*n* = 3 per species); statistical significance was assessed using one-way ANOVA followed by Tukey’s multiple-comparisons test. In the bottom panel, points represent individual cells. **c**, Projection of *ALCAM* expression onto the integrated force-directed layout of limb lateral plate mesoderm-derived cells, confirming enrichment within the PeriChon population (outlined). **d,** Cell-type-resolved chromatin accessibility at zooHAR.297 in developing human cranial skeletal tissue, showing open chromatin across chondrogenic and osteogenic lineages. Genomic coordinates are shown in the GRCh38 assembly. e,f, Cell-type annotations and *ALCAM* expression in developing human cranial chondrogenic populations (e, UMAP) and lineages undergoing intramembranous ossification (f, force-directed layout), illustrating coordinated *ALCAM* expression and zooHAR.297 accessibility in chondrogenic progenitors and suture mesenchymal cells.

**Supplementary Fig. 7.**
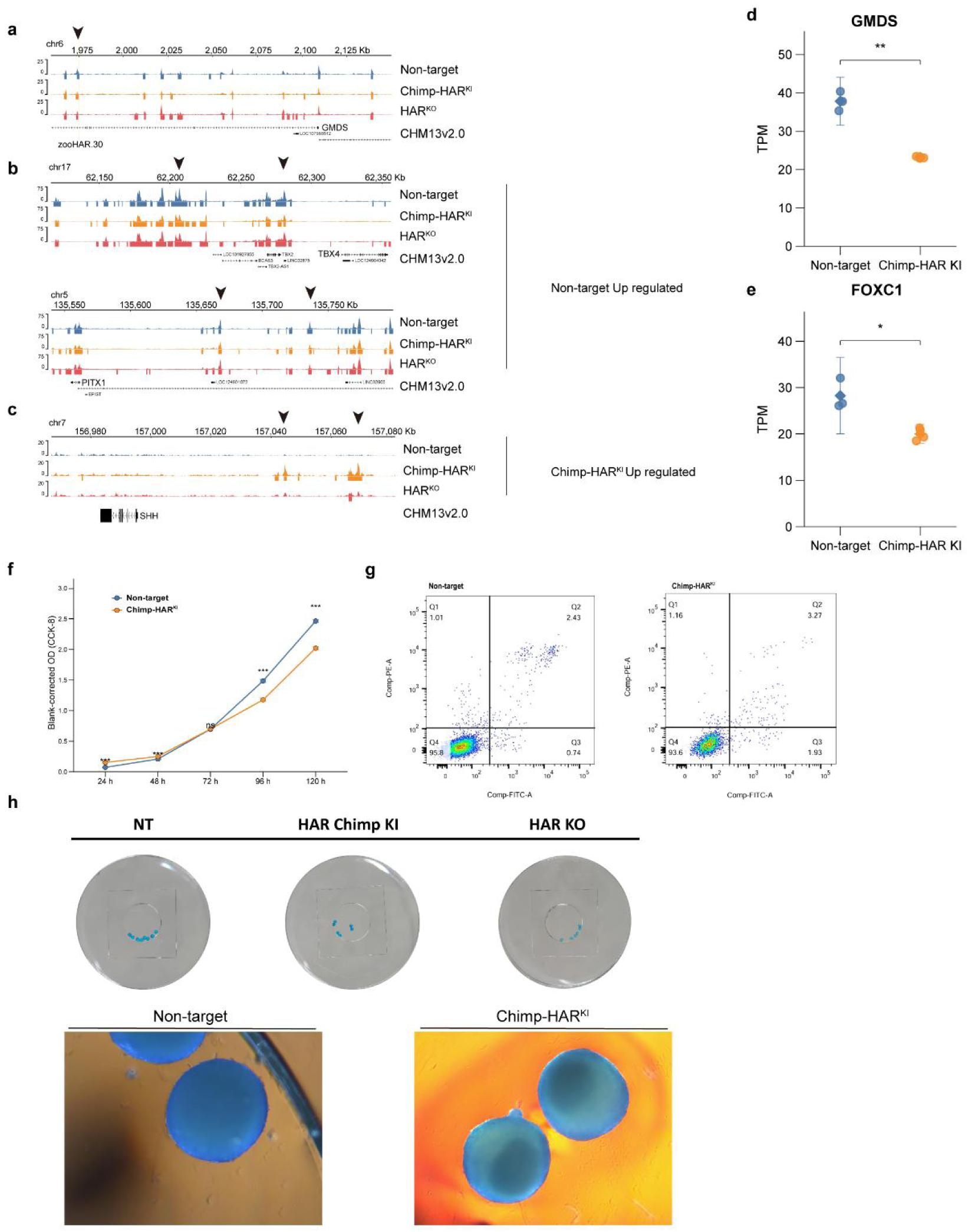
Chromatin and cellular effects of zooHAR.30 editing in human chondrocytes. **a–c,** H3K27ac CUT&Tag profiles in Non-target control, Chimp-HAR^KI^ and HAR^KO^ cells at the zooHAR.30–*GMDS* (**a**), *TBX4* and *PITX1* (**b**), and *SHH* (**c**) loci. Arrowheads indicate zooHAR.30 or representative regions with differential H3K27ac signal. Genomic coordinates are shown in the CHM13v2.0 assembly. **d,e,** Gene-level TPM expression of *GMDS*(**d**) and *FOXC1* (**e**) in Non-target and Chimp-HAR^KI^ cells. Circles represent individual RNA-seq samples (*n* = 3 Non-target and *n* = 4 Chimp-HAR^KI^); diamonds and error bars indicate mean ± 95% confidence interval. *P* values were calculated using two-sided Welch’s *t*-tests. **f,** Cell proliferation assessed by CCK-8 assay at the indicated time points. Points and error bars indicate mean ± s.e.m. Statistical significance was assessed using two-sided Welch’s *t*-tests separately at each time point. **g,** Representative flow cytometry plots of Annexin V and propidium iodide staining in Non-target and Chimp-HAR^KI^ cells. Numbers indicate the percentage of cells in each quadrant. **h,** Representative images of Alcian blue-stained three-dimensional chondrospheres from Non-target, Chimp-HAR^KI^ and HAR^KO^ cells after 21 days of chondrogenic induction at 5% O₂; enlarged images of Non-target and Chimp-HAR^KI^ chondrospheres are shown below. Images are representative of three independent experiments.

**Supplementary Fig. 8.**
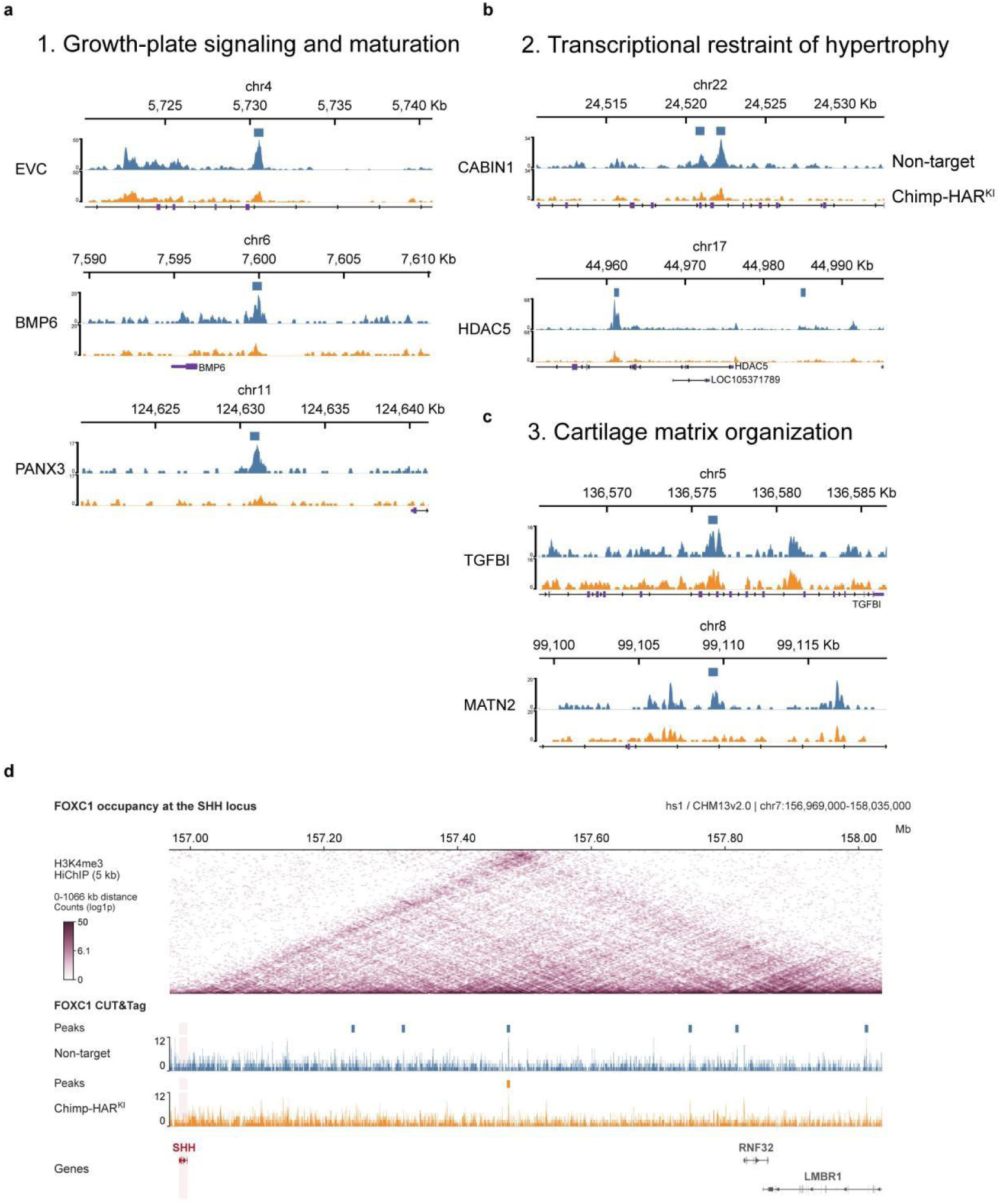
FOXC1 occupancy at osteochondral regulatory loci following zooHAR.30 sequence replacement. **a-c,** FOXC1 CUT&Tag profiles in Non-target and Chimp-HAR^KI^ cells at selected loci associated with growth-plate signalling and maturation (**a**), transcriptional regulation of hypertrophy (**b**) and cartilage matrix organization (**c**). Rectangles indicate FOXC1 peaks. **d,** H3K4me3 HiChIP contact map and FOXC1 CUT&Tag profiles at the *SHH* locus. Rectangles above the tracks indicate FOXC1 peaks in the corresponding cell group. Genomic coordinates are shown in the CHM13v2.0 assembly.

**Supplementary Fig. 9.**
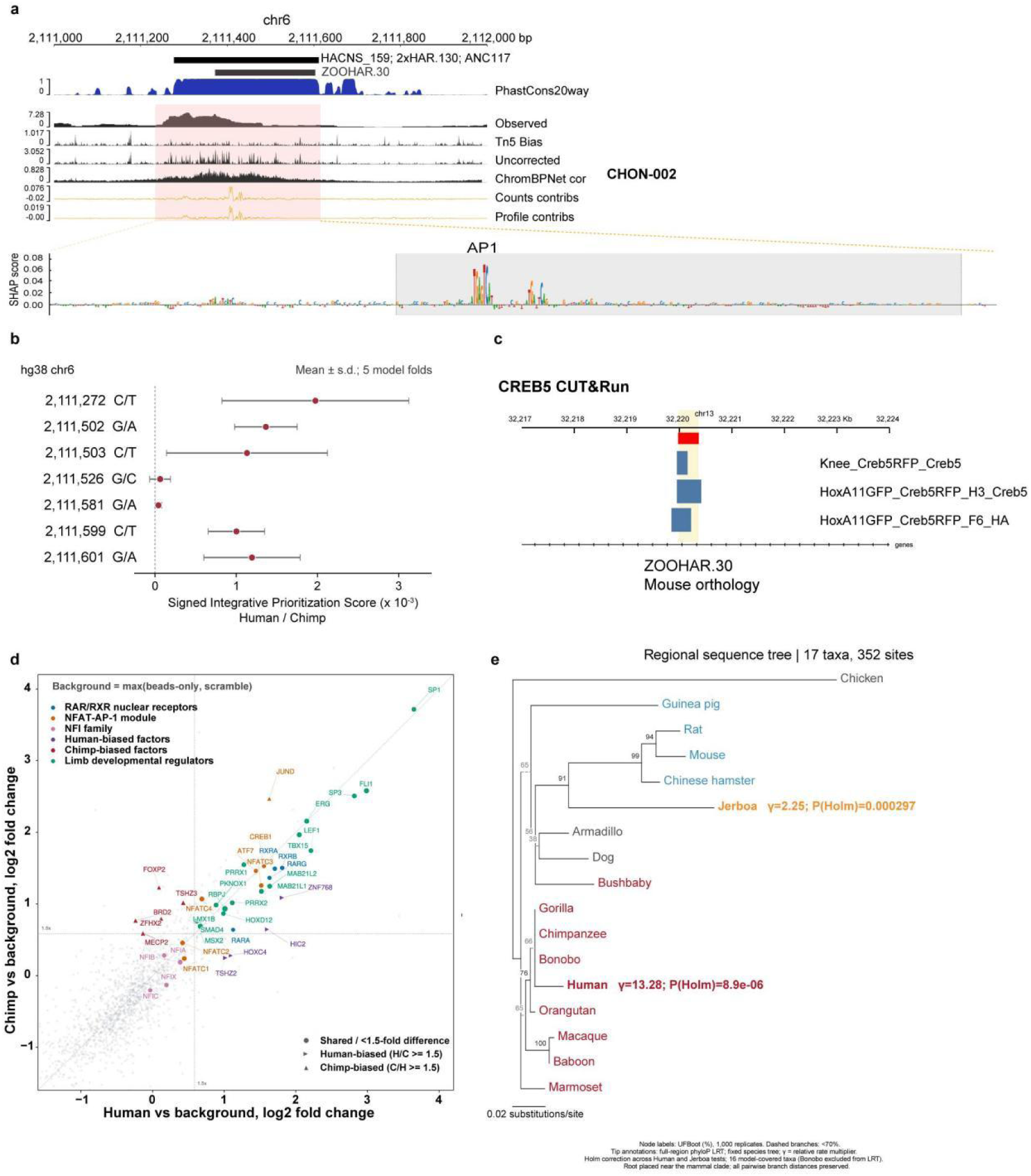
Sequence features and protein-binding landscape of zooHAR.30. **a,** Sequence conservation and ChromBPNet predictions at zooHAR.30 in CHON-002 cells, showing 20-way phastCons conservation, observed chromatin accessibility, predicted Tn5 bias, uncorrected and bias-corrected profiles, and base-level contribution scores for predicted counts and profiles. The enlarged contribution-score track highlights an AP-1 motif. **b,** ChromBPNet-predicted effects of seven individual human–chimpanzee nucleotide differences in the zooHAR.30-containing region, using a model trained on chromatin accessibility in human developing-limb InterZone cells. The signed integrative prioritization score (IPS) combines the predicted log₂ accessibility fold change for the human versus chimpanzee allele, Jensen–Shannon profile distance and active-allele quantile. Each substitution was evaluated individually in the hg38 sequence background. Points and error bars indicate mean ± s.d. across five model folds; nucleotide labels indicate the human and chimpanzee alleles, respectively. **c,** CREB5 CUT&RUN peaks at the mouse ortholog of zooHAR.30 in knee samples and in HOXA11-positive cells isolated from the developing zeugopod, as indicated. **d,** DNA pull-down mass spectrometry using human and chimpanzee zooHAR.30 sequences and mouse limb-bud nuclear extracts. Axes show log₂ fold enrichment over background, defined for each protein as the greater signal from the beads-only and scrambled-sequence controls. Point shapes indicate shared, human-biased or chimpanzee-biased proteins using a 1.5-fold sequence-comparison threshold; colors indicate selected protein groups. **e,** Regional sequence tree based on 352 aligned sites from 17 taxa. Node labels indicate ultrafast bootstrap support from 1,000 replicates; dashed branches indicate support below 70%. Human and jerboa tip annotations show relative rate multipliers (γ) and Holm-adjusted *P* values from full-region phyloP likelihood-ratio tests on a fixed species tree. Bonobo was excluded from these tests.

**Supplementary Fig. 10.**
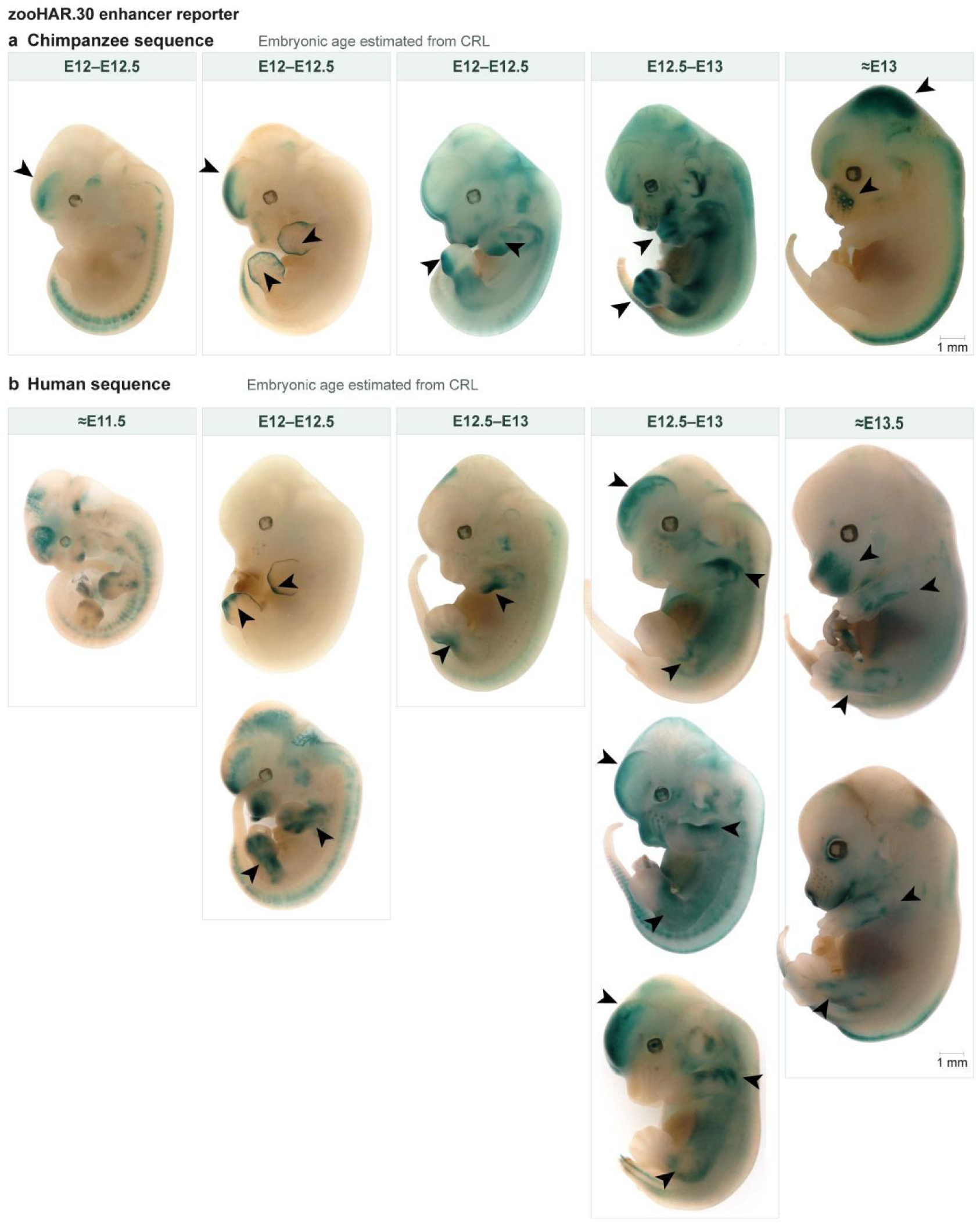
In vivo reporter activity of human and chimpanzee zooHAR.30 sequences. **a,b**, Whole-mount LacZ staining of transgenic mouse embryos carrying the chimpanzee (a) or human (b) zooHAR.30 sequence in a piggyBac-Hsp68-lacZ reporter. Approximate embryonic ages were estimated from crown-rump length (CRL), as indicated. Arrowheads highlight reporter activity in the head and developing limbs. Scale bars, 1 mm.

## References

1. Darwin, C. The Descent of Man, and Selection in Relation to Sex (John Murray, 1871).

2. Young, N. M., Wagner, G. P. & Hallgrímsson, B. Development and the evolvability of human limbs. Proc. Natl Acad. Sci. USA 107, 3400–3405 (2010).

3. Kun, E. et al. The genetic architecture and evolution of the human skeletal form. Science 381, eadf8009 (2023).

4. Carroll, S. B. Evolution at two levels: on genes and form. PLoS Biol. 3, e245 (2005).

5. Long, F. & Ornitz, D. M. Development of the endochondral skeleton. Cold Spring Harb. Perspect. Biol. 5, a008334 (2013).

6. Senevirathne, G. et al. The evolution of hominin bipedalism in two steps. Nature 645, 952–963 (2025).

7. To, K. et al. A multi-omic atlas of human embryonic skeletal development. Nature 635, 657–667 (2024).

8. Zhang, B. et al. A human embryonic limb cell atlas resolved in space and time. Nature 635, 668–678 (2024).

9. King, M. C. & Wilson, A. C. Evolution at two levels in humans and chimpanzees. Science 188, 107–116 (1975).

10. Prud’homme, B., Gompel, N. & Carroll, S. B. Emerging principles of regulatory evolution. Proc. Natl Acad. Sci. USA 104, 8605–8612 (2007).

11. Barr, K. A., Rhodes, K. L. & Gilad, Y. The relationship between regulatory changes in *cis* and *trans* and the evolution of gene expression in humans and chimpanzees. Genome Biol. 24, 207 (2023).

12. Cotney, J. et al. The evolution of lineage-specific regulatory activities in the human embryonic limb. Cell 154, 185–196 (2013).

13. Yan, Y. et al. The gene-regulatory evolution of the human skeleton. Nature 658, 698–706 (2026).

14. Okamoto, A. S., Coveney, C. R., Ganapathee, D. S. & Capellini, T. D. *In vitro* massively parallel screening of human regulatory elements involved in postcranial skeletal development for differential activity compared to chimpanzee. Genome Biol. Evol. 18, evag121 (2026).

15. Pollard, K. S. et al. Forces shaping the fastest evolving regions in the human genome. PLoS Genet. 2, e168 (2006).

16. Prabhakar, S., Noonan, J. P., Pääbo, S. & Rubin, E. M. Accelerated evolution of conserved noncoding sequences in humans. Science 314, 786 (2006).

17. Bird, C. P. et al. Fast-evolving noncoding sequences in the human genome. Genome Biol. 8, R118 (2007).

18. Bush, E. C. & Lahn, B. T. A genome-wide screen for noncoding elements important in primate evolution. BMC Evol. Biol. 8, 17 (2008).

19. Keough, K. C. et al. Three-dimensional genome rewiring in loci with human accelerated regions. Science 380, eabm1696 (2023).

20. Capra, J. A., Erwin, G. D., McKinsey, G., Rubenstein, J. L. R. & Pollard, K. S. Many human accelerated regions are developmental enhancers. Philos. Trans. R. Soc. Lond. B Biol. Sci. 368, 20130025 (2013).

21. Dutrow, E. V. et al. Modeling uniquely human gene regulatory function via targeted humanization of the mouse genome. Nat. Commun. 13, 304 (2022).

22. Prabhakar, S. et al. Human-specific gain of function in a developmental enhancer. Science 321, 1346–1350 (2008).

23. Pal, A. et al. Resolving the three-dimensional interactome of human accelerated regions during human and chimpanzee neurodevelopment. Cell 188, 1504–1523.e27 (2025).

24. Schindler, M. et al. Comparative single-cell analyses reveal evolutionary repurposing of a conserved gene programme in bat wing development. *Nat*. Ecol. Evol. 9, 1626–1642 (2025).

25. He, P. et al. The changing mouse embryo transcriptome at whole tissue and single-cell resolution. Nature 583, 760–767 (2020).

26. Hrovatin, K. et al. Integrating single-cell RNA-seq datasets with substantial batch effects. BMC Genomics 26, 974 (2025).

27. Xue, J. R. et al. The functional and evolutionary impacts of human-specific deletions in conserved elements. Science 380, eabn2253 (2023).

28. Andrey, G. et al. A switch between topological domains underlies *HoxD* genes collinearity in mouse limbs. Science 340, 1234167 (2013).

29. Narendra, V., Bulajić, M., Dekker, J., Mazzoni, E. O. & Reinberg, D. CTCF-mediated topological boundaries during development foster appropriate gene regulation. Genes Dev. 30, 2657–2662 (2016).

30. Kragesteen, B. K. et al. Dynamic 3D chromatin architecture contributes to enhancer specificity and limb morphogenesis. Nat. Genet. 50, 1463–1473 (2018).

31. Sun, J., Ishii, M., Ting, M.-C. & Maxson, R. *Foxc1* controls the growth of the murine frontal bone rudiment by direct regulation of a Bmp response threshold of *Msx2*. Development 140, 1034–1044 (2013).

32. Yoshida, M. et al. The transcription factor Foxc1 is necessary for Ihh–Gli2-regulated endochondral ossification. Nat. Commun. 6, 6653 (2015).

33. Rice, R., Rice, D. P. C., Olsen, B. R. & Thesleff, I. Progression of calvarial bone development requires *Foxc1* regulation of *Msx2* and *Alx4*. Dev. Biol. 262, 75–87 (2003).

34. Arai, F., Ohneda, O., Miyamoto, T., Zhang, X. Q. & Suda, T. Mesenchymal stem cells in perichondrium express activated leukocyte cell adhesion molecule and participate in bone marrow formation. J. Exp. Med. 195, 1549–1563 (2002).

35. Sekine, K. et al. *Fgf10* is essential for limb and lung formation. Nat. Genet. 21, 138–141 (1999).

36. Logan, M. & Tabin, C. J. Role of *Pitx1* upstream of *Tbx4* in specification of hindlimb identity. Science 283, 1736–1739 (1999).

37. Duboc, V. et al. *Tbx4* function during hindlimb development reveals a mechanism that explains the origins of proximal limb defects. Development 148, dev199580 (2021).

38. Zhu, J. et al. Uncoupling Sonic hedgehog control of pattern and expansion of the developing limb bud. Dev. Cell 14, 624–632 (2008).

39. Riddle, R. D., Johnson, R. L., Laufer, E. & Tabin, C. Sonic hedgehog mediates the polarizing activity of the ZPA. Cell 75, 1401–1416 (1993).

40. Grimsrud, C. D. et al. BMP-6 is an autocrine stimulator of chondrocyte differentiation. J. Bone Miner. Res. 14, 475–482 (1999).

41. Iwamoto, T. et al. Pannexin 3 regulates intracellular ATP/cAMP levels and promotes chondrocyte differentiation. J. Biol. Chem. 285, 18948–18958 (2010).

42. Nishimori, S., et al. PTHrP targets HDAC4 and HDAC5 to repress chondrocyte hypertrophy. JCI Insight 4, e97903 (2019).

43. Lee, J.-M., Lee, E.-H., Kim, I.-S. & Kim, J.-E. *Tgfbi* deficiency leads to a reduction in skeletal size and degradation of the bone matrix. Calcif. Tissue Int. 96, 56–64 (2015).

44. Richard, D. et al. Evolutionary selection and constraint on human knee chondrocyte regulation impacts osteoarthritis risk. Cell 181, 362–381.e28 (2020).

45. Conery, M. et al. GWAS-informed data integration and non-coding CRISPRi screen illuminate genetic etiology of bone mineral density. Genome Biol. 26, 331 (2025).

46. Hatzikotoulas, K. et al. Translational genomics of osteoarthritis in 1,962,069 individuals. Nature 641, 1217–1224 (2025).

47. Bartell, E. et al. Genetics of skeletal proportions across two different populations. Am. J. Hum. Genet. 113, 794–808 (2026).

48. Chen, H. et al. Heads, shoulders, elbows, knees, and toes: modular *Gdf5* enhancers control different joints in the vertebrate skeleton. PLoS Genet. 12, e1006454 (2016).

49. Muthuirulan, P. et al. Joint disease-specificity at the regulatory base-pair level. Nat. Commun. 12, 4161 (2021).

50. Pampari, A. et al. ChromBPNet: bias factorized, base-resolution deep learning models of chromatin accessibility reveal *cis*-regulatory sequence syntax, transcription factor footprints and regulatory variants. Preprint at bioRxiv 10.1101/2024.12.25.630221 (2024).

51. Oldfield, A. J. et al. Histone-fold domain protein NF-Y promotes chromatin accessibility for cell type-specific master transcription factors. Mol. Cell 55, 708–722 (2014).

52. Zhang, C.-H. et al. *Creb5* coordinates synovial joint formation with the genesis of articular cartilage. Nat. Commun. 13, 7295 (2022).

53. Zhang, C.-H., et al. Creb5 controls its own expression and directly induces the joint interzone regulatory program. Proc. Natl Acad. Sci. USA 122, e2501830122 (2025).

54. Haack, H. & Gruss, P. The establishment of murine *Hox-1* expression domains during patterning of the limb. Dev. Biol. 157, 410–422 (1993).

55. Mercader, N. et al. Ectopic *Meis1* expression in the mouse limb bud alters P-D patterning in a *Pbx1*-independent manner. Int. J. Dev. Biol. 53, 1483–1494 (2009).

56. Holliday, T. W. et al. Body size and proportions of *Australopithecus sediba*. PaleoAnthropology 2018, 406–422 (2018).

57. Lay, K., Kume, T. & Fuchs, E. FOXC1 maintains the hair follicle stem cell niche and governs stem cell quiescence to preserve long-term tissue-regenerating potential. Proc. Natl Acad. Sci. USA 113, E1506–E1515 (2016).

58. Wang, L., Siegenthaler, J. A., Dowell, R. D. & Yi, R. Foxc1 reinforces quiescence in self-renewing hair follicle stem cells. Science 351, 613–617 (2016).

59. Wang, Y. Construction of 3D Genome Regulatory Maps of the Lower Limb Chondrocytes of Human and Rhesus Macaque. Master’s thesis, University of Chinese Academy of Sciences (2021).

60. Mumbach, M. R. et al. HiChIP: efficient and sensitive analysis of protein-directed genome architecture. Nat. Methods 13, 919–922 (2016).

61. Blanco, M. J. et al. Tracing gene expression through detection of β-galactosidase activity in whole mouse embryos. J. Vis. Exp. 136, e57785 (2018).

